# Object boundary detection in natural images may depend on ‘incitatory’ cell-cell interactions

**DOI:** 10.1101/436949

**Authors:** Gabriel C. Mel, Chaithanya A. Ramachandra, Bartlett W. Mel

## Abstract

Detecting object boundaries is crucial for recognition, but how the process unfolds in visual cortex remains unknown. To study the problem faced by a hypothetical boundary cell, and to predict how cortical circuitry could produce a boundary cell from a population of conventional “simple cells”, we labeled 30,000 natural image patches and used Bayes’ rule to help determine how a simple cell should influence a nearby boundary cell depending on its relative offset in receptive field position and orientation. We identified three basic types of cell-cell interactions: rising and falling interactions with a range of slopes and saturation rates, as well as non-monotonic (bump-shaped) interactions with varying modes and amplitudes. Using simple models we show that a ubiquitous cortical circuit motif consisting of direct excitation and indirect inhibition – a compound effect we call “incitation” – can produce the entire spectrum of simple cell-boundary cell interactions found in our dataset. Moreover, we show that the synaptic weights that parameterize an incitation circuit can be learned by a single-layer “delta” rule. We conclude that incitatory interconnections are a generally useful computing mechanism that the cortex may exploit to help solve difficult natural classification problems.

**Significance statement:** Simple cells in primary visual cortex (V1) respond to oriented edges, and have long been supposed to detect object boundaries, yet the prevailing model of a simple cell – a divisively normalized linear filter – is a surprisingly poor natural boundary detector. To understand why, we analyzed image statistics on and off object boundaries, allowing us to characterize the neural-style computations needed to perform well at this difficult natural classification task. We show that a simple circuit motif known to exist in V1 is capable of extracting high-quality boundary probability signals from local populations of simple cells. Our findings suggest a new, more general way of conceptualizing cell-cell interconnections in the cortex.

## Introduction

The primary visual cortex (area V1) is a complex, poorly understood, multi-purpose image processor optimized to extract information from natural scenes – which are themselves complex, poorly understood signals. Thus, understanding how V1 operates presents a challenging reverse engineering problem. A longstanding hypothesis is that orientation-tuned V1 cells somehow participate in object boundary detection, a core process in biological vision (Biederman, 1987; Gilbert and Wiesel, 1990; Heydt and Peterhans, 1989; Hubel and Wiesel, 1962; Kapadia et al., 1995) that is crucial for the functions of both ventral and dorsal streams (Biederman, 1987; Hoffman, 2000; Rust and Dicarlo, 2010; Theys et al., 2015). However, little progress has been made in refining or testing this hypothesis, in part due to our lack of understanding of the structure of natural object boundaries, and particularly, what a V1 cell needs to do to reliably distinguish boundaries from non-boundaries.

This uncertainty has made it difficult to form specific computational hypotheses as to how V1 circuits perform this behaviorally-relevant classification task. Previous work has analyzed natural image statistics to determine how local boundary segments are arranged in images (Sanguinetti et al., 2010; Sigman et al., 2001), and how these arrangements relate to human contour grouping performance (W. S. Geisler, Perry, Super, & Gallogly, 2001). However, no study has yet attempted to deconstruct the natural boundary *detection* problem in detail, or to link the computations necessary for boundary detection to particular neural mechanisms.

With the goal to better understand the computations underlying object boundary detection in V1 (Figure 1A), we began with a known cell type – orientation-tuned “simple cells” (as defined by Hubel and Wiesel, 1962), and typically modeled as divisively normalized oriented linear filters (Carandini and Heeger, 2012) – and asked how the outputs of a population of simple cells (SCs), whose receptive fields (RFs) densely cover an area of the visual field, should be combined to produce a “boundary cell” (BC) whose firing rate represents the probability that an object boundary is present within its RF (Figure 1B). When framed in this way, Bayes’ rule tells us what data to extract from natural images to obtain an answer to the question. In a previous study (Ramachandra and Mel, 2013), we noted that under the simplifying assumption of “class conditional independence” (see methods for a detailed discussion), simple cell-boundary cell interactions are captured by the log-likelihood ratio (LLR) functions embedded in Bayes’ rule (colored expressions in Figure 1C), which represent the evidence that a given simple cell provides about the presence of an object boundary within a neighboring boundary cell’s receptive field (Figure 1D). We found that SC-BC interactions were diverse, and in some cases involved compound excitatory and inhibitory effects. However, since only a small number of cells was analyzed in that study, we could not come to general conclusions about the types of cell-cell interactions needed to compute boundary probability, making it difficult to compare and contrast possible neural mechanisms.

**Figure 1.**
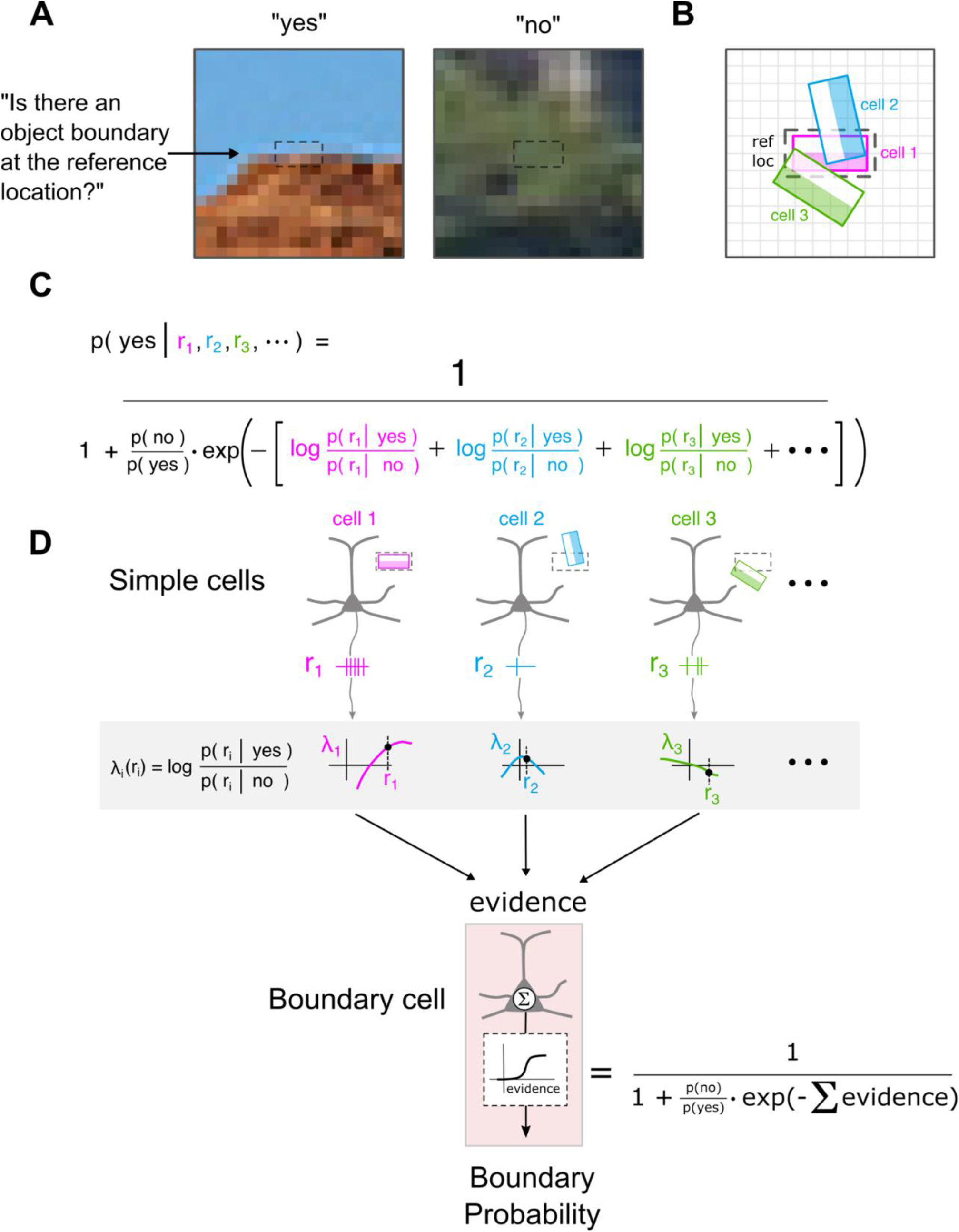
Calculating boundary probability from natural images using Bayes’ rule. **A**, The boundary detection problem can be encapsulated by the question and answers shown; ∼30,000 natural image patches were classified in this way. Dashed box indicates a “reference location” where a boundary might appear. Patches shown during labelling were 20×20 pixels. **B**, 3 (of 300) oriented cells with responses r_1_, r_2_, r_3_ are shown in the vicinity of the reference location. **C,** Under the assumption that simple cell responses are class-conditionally independent (see main text), Bayes’ rule gives an expression for boundary probability in terms of individual cell log-likelihood ratios (LLRs) (colored terms in denominator). **D**, Cell responses *r*_*i*_ are passed through their respective LLR functions *λ*_*i*_(), and the results are summed and passed through a fixed sigmoidal “f-I curve” to yield boundary probability.

In this study, we analyze a much larger dataset, and compute the full set of simple cell-boundary cell interaction functions for a population of 300 simple cells surrounding a “reference location” where a boundary might be detected. We find that the simple cell-boundary cell interactions suggested by the natural image LLR functions follow a predictable pattern that depends on the offset in position and orientation between simple cell and boundary cell receptive fields, and we show that a well-known cortical circuit motif can implement the entire spectrum of individual SC-BC interactions found in our data set. Finally, we demonstrate that a cortically-inspired neural network can produce a boundary-detecting cell from simple cells with a single layer of excitatory synapses and a single inhibitory interneuron. Our findings suggest that cortical sensory computations, including the detection of natural object boundaries, may depend on a specific class of structured excitatory-inhibitory cell-cell interactions.

## Materials and methods

### Image preprocessing

As in Ramachandra and Mel (2012), we used a modified version of the COREL database for boundary labeling in natural images. Several image categories, including sunsets and paintings were removed from the full COREL database since their boundary statistics differed markedly from that of typical natural images. Custom code was used to select ∼30,000 20×20 pixel image patches for labelling. The “reference location” representing a hypothetical boundary cell’s receptive field location was defined as the elongated, horizontal 2×4 pixel region at the center of the patch (dashed box, Figure 1A, B).

### Natural image data collection

To collect ground-truth data relating to natural contour statistics, for each image patch to be labeled, a horizontal 2×4 pixel rectangular box was drawn around a centered “reference location” and human labelers were asked to answer the question, “On a scale from 1 to 5, with 1 meaning ‘extremely unlikely’ and 5 meaning ‘extremely likely’ – how likely is it that there is an object boundary passing horizontally through the reference box, end to end, without leaving the box?” To qualify as valid, boundary segments also had to be visible and unoccluded within the box. We restricted labelling to horizontal boundaries (i.e. horizontal reference boxes) since pixel lattice discretization made it more difficult to judge oblique orientations, and because we expected cell response statistics in natural images to be roughly orientation invariant. (This expectation was supported by subsequent tests showing that LLR functions obtained for horizontal boundaries also led to high boundary detection performance on oblique boundaries). Labeler responses were recorded, and patches with scores of 1 or 2, were classified as “no” patches, while patches with scores of 4 or 5 were classified as “yes” patches. Agreement between labelers was very high, based on informal observations when two labelers worked together. Rare ambiguous patches that could cause labeler disagreement were often given scores of 3, so these patches were excluded from our analyses. After labeling, the dataset was doubled by adding left-right flipped versions of each patch, and assigning the same label as the un-flipped counterpart.

### Collecting virtual simple cell responses on and off natural boundaries

Given a large set of image patches, some labeled as boundaries and others not, the next step was to collect virtual simple cell responses densely covering boundary vs. non-boundary image patches so that their different statistics could be analyzed. Original color image patches were converted to single-channel (monochrome) intensity images (0.29 *R* + 0.59 *G* + 0.11 *B*). Simple cell-like oriented linear “filters” were created by rotating a 2×4 pixel horizontal filter kernel in 15° increments from 0 to 165° (i.e. 12 orientations). The filter kernel for the horizontal (0°) cell is shown in Figure 2A. Positive filter coefficients represent ON subregions, and negative coefficients represent OFF subregions of the virtual simple cell’s receptive field. Blank kernel entries represent zeros. Filter coefficients in rotated kernels were computed by rotating the horizontal kernel using “bilinear interpolation” (https://en.wikipedia.org/wiki/Bilinear_interpolation) (Figure 2B, C). In essence, any entry in an oriented kernel that was affected by the rotated horizontal pattern would be assigned a coefficient interpolated based the nearest 4 coefficients in the original horizontal pattern after back-rotation. (The locations of the faint blue dots in Figure 2A show the locations of the dark blue dots in Figure 2B rotated back by 45° into the horizontal frame of references; the 4 coefficients closest to each faint blue dot were used for the interpolation). A similar procedure was used to generate all other rotated filter kernels.

**Figure 2.**
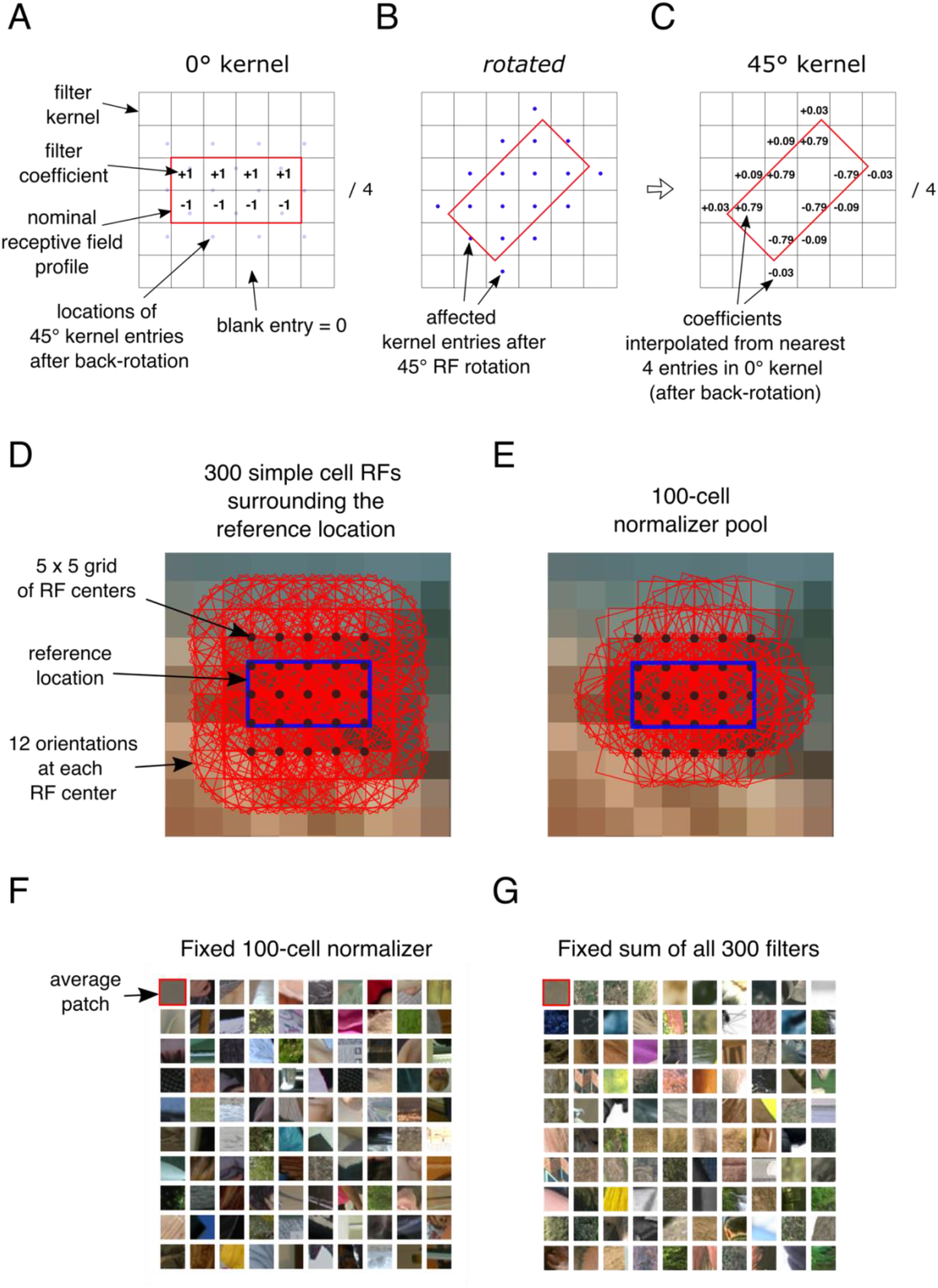
Simple cell “filter” construction, layout, and normalization pools. **A,** Diagram shows layout of positive and negative filter coefficients for computing simple cells responses. Positive coefficients represent the ON subregion of the receptive field (RF), and negative coefficients represent the OFF subregion. Faint blue dots indicate off-grid positions (in x-y space) of the 18 pixel locations holding coefficients for the 45° case shown in panel B, back rotated to the horizontal orientation. **B,** Blue dots indicate all pixels that overlap with the red box, representing the original horizontal receptive field rotated by 45°. Each of these 18 pixels will contain a coefficient (see panel C), computed from bilinear interpolation of the coefficients in the original horizontal layout (see panel A). The coefficients “down the middle” of the 45° filter are zero, reflecting an equal contribution from the ON and OFF subregions of the RF. **D,** Layout of 300 filter locations (red) in 5×5 grid surrounding the reference location (blue). **E,** Subset of 100 of the 300 simple cell RFs chosen as the normalizer pool in the following figures. See text for details. **F,** Examples of image patches drawn at random from the image database whose normalizer values (i.e. the sum of the response magnitudes of the 100 cells shown in E) fell into the same bin (200 +/- 40); see text for details. Upper left patch is the average of all. **G,** Same as panel F, but with patch selection based on a different normalizer: the sum of the response magnitudes of all 300 simple cells covering the region (whose RFs are shown in panel D).

A simple cell’s “response” value at a particular image location was computed as the dot product between the cell’s filter kernel and the underlying image intensity pixel values. A simple cell generated a positive response when its positive kernel coefficients mostly overlapped with bright image pixels and its negative coefficients mostly overlapped with dark image pixels. The largest positive responses occurred on light-dark boundaries of a cell’s preferred orientation. A negative response was treated as a positive response of a distinct simple cell with opposite contrast polarity (i.e. rotated by 180°).

Given a labeled image patch, simple cell responses covering that patch were collected on a 5 x 5 grid centered on the patch, at each of 12 orientations (Figure 2D). This led to 5 x 5 x 12 = 300 simple cells responding to any given image patch. Simple cell response data was accumulated separately for “yes” and “no” labeled patches.

### Restriction to “normalized” image patches

The prevailing model of a simple cell consists of linear filter whose output is divisively normalized by activity in the cell’s surround (Carandini and Heeger, 2012). Normalizing a simple cell’s response involves (1) computing the cell’s pre-normalized response (e.g., as we do above using an oriented linear filter); (2) measuring the sum N of other neural activity in the cell’s surround including cells of the same or different orientation located in the near or far surround; and (3) inhibiting the simple cell’s response as an increasing function of N. Normalization is often called “divisive” as the surround activity term, N, generally appears in the denominator of the overall cell response expression. Response normalization plays a variety of roles in the brain (see Carandini and Heeger, 2012 for an excellent review), though in the visual system normalization is most often discussed as a means of counteracting the effect of multiplicative “nuisance” factors that modulate the responses of entire local population of neurons in a correlated fashion. For example, if the level of illumination varies across the visual field, a common occurrence in natural scenes, the spatial variation in light level tends to drive up response rates of all neurons in brighter areas and drive down response rates of all neurons in darker areas. Normalization circuitry helps to cancel out these correlated, population-level neural response variations, leading to a greater degree of illumination invariance across the visual field, and a greater degree of statistical independence in the neural code (Wainwright & Simoncelli, 2000; Schwartz & Simoncelli, 2001; Liang, Simoncelli, & Lei, 2000; Zetsche & Rohrbein, 2001; Parra, Spence, & Sajda, 2000; Fine et al., 2003; Karklin & Lewicki, 2003, 2005; Zhou and Mel 2008).

The importance of normalization with respect to our analysis is that neural activity in the non-classical surround of a simple cell can powerfully influence its response, and given that our study involves collecting and statistically analyzing simple cell responses in natural images, we made efforts to take account of the effects of normalization circuitry on simple cell responses. As discussed above, the most common assumption about response normalization in the cortex is that it depends on the summed activity of a pool of neurons in the vicinity of a target cell (see Carandini and Heeger 2012 for review), and that the normalization function is “divisive” (meaning the normalization term appears in the denominator). We adopted this assumption, and selected a pool of 100 oriented cells from the 300 shown in Figure 2D using an ad-hoc procedure designed to find a subset of the cells that would be minimally correlated in normalized patches. (Limiting the normalizing pool to a subset of the 300 cells turned out not to be unnecessary, as we found that including all 300 cells in the normalizer pool led to functionally equivalent results, but we describe the selection procedure below since the 100-cell normalizer was what was actually used to generate our result figures. The reader not interested in the details of the normalization approach can skip to the next paragraph). We started by defining a “basic normalizer” consisting of the single linear filter value at the reference location. We then culled out a large set of natural image patches whose basic normalizer values fell into a narrow range, that is, all the culled patches had roughly the same filter value at the reference location. The value used for the basic normalizer was 10±2, but the particular value mattered little; what mattered was that the value was fixed across all image patches collected. We then incrementally “grew” a normalization pool as follows. The cell at the reference location was considered to be the first cell in the normalization pool (C_1_). A second cell, call it C_2_, was added to the pool by choosing that cell (from the 299 remaining) for which the correlation of its absolute value to that of the other cells in the pool (which was only C_1_ in this case) was closest to zero. The absolute value was used because negative values were considered to be responses of distinct cells with opposite polarity. A third cell C_3_ was then chosen (from the 298 remaining) for which the correlation of its absolute value to that of C_1_ and C_2_ was, on average, closest to zero. This “greedy” (”choose the best each time”) procedure continued until 100 cells were chosen, leading to the particular subset of cell RFs shown in Figure 2E. The set of 100 cells is also specified in the Matlab file “normalizer_filter_subs.mat” in the Supplementary Materials. Having chosen the 100-cell normalization pool, we could now cull out a second-generation-normalized set of image patches from the image database by selecting patches for which the sum of the responses of all 100 cells in the normalization pool (converted to absolute values) fell into a fixed response range N=200±40. The data in our results figures was generated from this set of normalized image patches. However, as mentioned above, random image patches normalized using all 300 filters surrounding the reference location (and N≈600 +/- 40; exact bin center chosen to match mean on our normalized labelled patch database) were indistinguishable as a group (see examples in Figures 2F and 2G), had indistinguishable averages (upper left each image grid), and led to LLR functions for boundary detection that were very similar.

The restriction of our analysis to image patches within a narrow range of normalizer values was functionally similar to carrying out our analysis “under controlled lighting conditions”. The benefit of this restriction was that it allowed us to probe the relationships between simple cell responses and boundary probability without having to assume that we know the precise form of the function that normalizes neural responses against changes in lighting or other regional correlating factors. The cost of this restriction is that we obtain information only about the boundary probability computation at a specific normalization level. This would be a serious problem if the boundary probability computation changed significantly at different levels of normalization. To mitigate this risk we verified that our analysis produced similar main results for 2 other value ranges of the 100-cell normalizer (N = 300 ± 40 and N = 400 ± 40). We accomplished this by assuming that the normalizing function divisively scales filter responses, which, given that the filters are linear, is equivalent to divisively scaling the image patches themselves. When we scaled down the image patches from the other two normalizer bins by factors of 300/200 = 1.5 and 400/200 = 2, respectively, and thereafter processed all image patches equivalently, we obtained results that were functionally indistinguishable from those produced from the original “200” dataset (data not shown). In our main results figures, we include only the data derived from image patches in the original “200” normalizer bin.

### Bayesian formalism

We assume that a boundary cell computes *p*( *yes* | *r*_1_ *r*_2_ *r*_3_ …), that is, the probability that a boundary is present at the reference location given a set of simple cell responses *r*_1_ *r*_2_ *r*_3_, etc., whose receptive fields surround and cover the reference location (Fig. 1D). Using Bayes’ rule we obtain

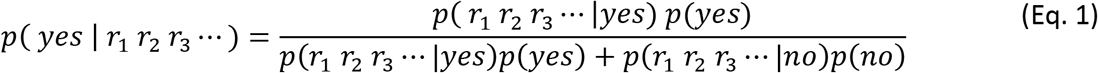

Dividing through by the numerator and rearranging, we find

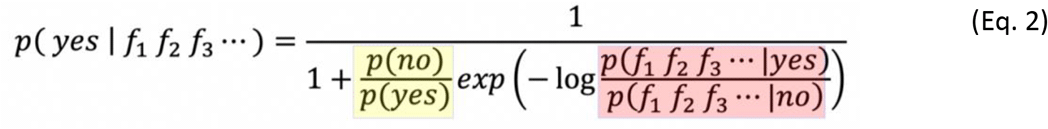

The log term in the denominator (shaded pink) of Eq. 2 consists of a ratio of two “likelihoods”. The numerator and denominator of the likelihood ratio represent, respectively, the probability of measuring a particular combination of simple cell responses (*r*_1_ *r*_2_ *r*_3_ …) when a boundary is present at the reference location (*yes*), and the probability of measuring the *same* combination of filter responses when a boundary is not present at the reference location (*no*). When a simple cell response combination (*r*_1_ *r*_2_ *r*_3_ …) is measured in the vicinity of a reference location, and based on statistical record keeping we know that that response combination is more likely to be associated with a *yes* case than a *no* case, then the likelihood ratio is greater than 1, meaning there is positive evidence for a boundary; the logarithm term is greater than 0, and the exponential term drops below 1 due to the negative sign of the exponent. When the likelihood ratio is *much* greater than 1, the exponential term approaches zero, and the denominator of the full expression approaches 1, indicating that a boundary at the reference location is highly probable. The yellow-shaded term in the denominator of Eq. 2 is the prior “odds” of finding an object boundary at a randomly sampled location, which is the ratio of the prior probabilities of “yes” and “no” cases. Yes cases made up 2.4% of our default dataset (for N=200), meaning the odds of a boundary was 2.4/97.6 = 2.45%. The odds term functions as a sort of evidence threshold: the lower the prior odds of a boundary, the stronger the evidence must be in order to reach a 50% probability that a boundary is present at the reference location. For prior odds of 2.45%, a likelihood ratio of 41 would be needed to reach a 50% boundary probability.

### Simplifying Bayes rule by assuming class conditional independence (CCI)

Using Bayes rule as a tool for interpreting cortical circuit interactions runs into the obstacle that the likelihood expressions contributing to the likelihood ratio involve high-dimensional probability densities, where the dimension corresponds to the number of simple cells that will contribute to the boundary probability calculation (which could number in the 100’s). Collecting high-dimensional probability density functions from natural images, and representing them whether mathematically, by rote tabulation, or neurally, is for all intents and purposes intractable. However, if we assume that neighboring simple cells display a certain kind of statistical independence – i.e. “class conditional” independence – this radically simplifies the computational problem by allowing the high-dimensional likelihoods in Bayes rule to be factored and re-expressed as a sum of 1-dimensional log likelihood ratio (LLR) functions *λ*_*i*_(*r*_*i*_). These 1-dimensional LLR functions can be interpreted as “the varying effect that cell *i* has on a nearby boundary cell over its range of responses *r*_*i*_”. Fortuitously, the functions *λ*_*i*_ are simple in form (i.e. smooth and unimodal), and can be easily computed by a known cortical circuit motif. We therefore begin our analysis under the CCI assumption, as it provides a stepping stone to understanding the types of cell-cell interactions we should expect to find in a cortical circuit in which boundary probability is computed. We then take steps to bridge the gap between the predictions of the simplified analysis that assumes class conditional independence, and the types of cell-cell interactions that we should expect to find in the more realistic scenario where the simple cells contributing to a boundary probability computation do not satisfy the CCI assumption.

The assumption of class-conditional independence in our scenario implies that simple cell responses are statistically independent both for the set of image patches where a boundary is present at the reference location, and for the set of image patches where a boundary is not present at the reference location. When CCI holds, both the numerator and denominator terms in the likelihood ratio of Eq. 2 can be factored into individual cell response terms, and converted to a sum of cell-specific log likelihood ratio (LLRs) functions, as follows:

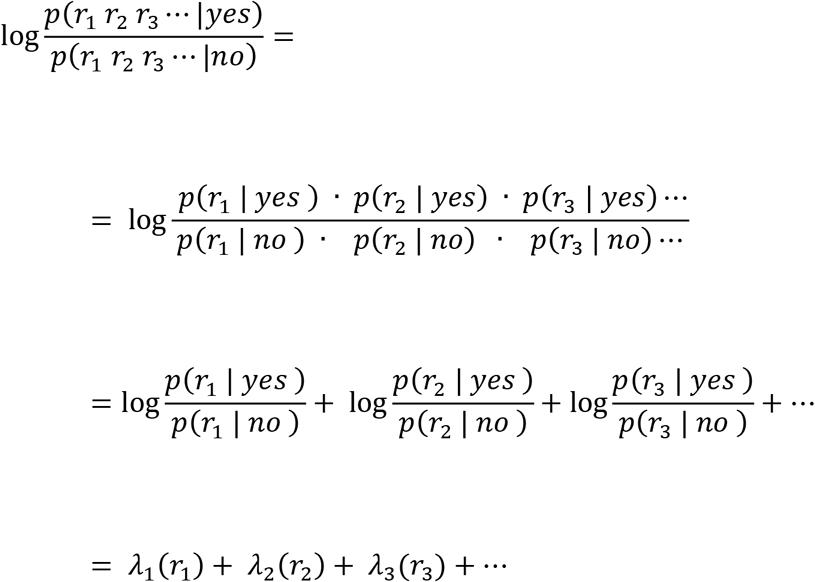

where *λ*_*i*_(*r*_*i*_) is the log likelihood ratio function for the *i*^*th*^ neighboring cell. In simple terms, the LLR function *λ*_*i*_(*r*_*i*_) captures how the evidence for a boundary at the reference location depends on the response of the *i*^*th*^ neighboring simple cell. We can write the CCI version of Bayes rule as

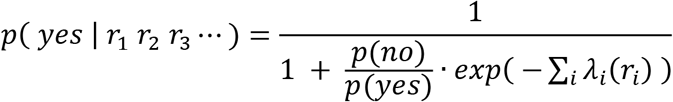

which reduces to the remarkably simple formula

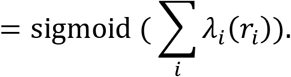

In intuitive terms, this equation says that for a boundary cell at the reference location to compute boundary probability within its receptive field, the response of each neighboring simple cell *r*_*i*_ should be passed through a nonlinear function *λ*_*i*_ whose form depends on the neighbor’s offset in position and orientation from the reference location; the results from all contributing simple cells should be summed; and the total should be passed through a final sigmoidal nonlinearity to produce the boundary cell’s response (Figure 1).

### Extracting the log likelihood ratio functions *λ*_*i*_(*r*_*i*_) from natural images

Histograms of the responses of the 300 simple cells surrounding the reference location were collected separately for “yes” and “no” image patches (total of 600 histograms). Histograms for no patches contained 50 evenly spaced bins. Yes histograms were binned more coarsely because our dataset had many fewer yes patches than no patches; between 8 and 20 evenly spaced bins were used to ensure smoothness of the response histograms for all cells. The yes and no histograms for each simple cell were converted to yes and no probability density functions (pdfs) by dividing the sample count in each bin by two factors: (1) the total sample count in the respective histogram, and (2) the bin width. From these pdfs, LLR functions λ(*r*) were constructed for each simple cell as 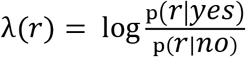, where *p*(*r*|*yes*) and *p*(*r*|*no*) were the cell’s yes and no pdf bin values, respectively. The log function was the natural logarithm (base e). To control noise levels, each cell’s λ function was considered valid only at non-extreme response levels for which the probability in both the yes and no pdfs exceeded minimum thresholds ( *p*(*r*|*yes*) > 0.005 and *p*(*r*|*no*) > 0.002). Different thresholds were used because smaller probabilities could be estimated more reliably in the no histograms given the much larger set of no image patches. Only data inside valid response regions is plotted in Figures 3-5 and Figure 7.

**Figure 3.**
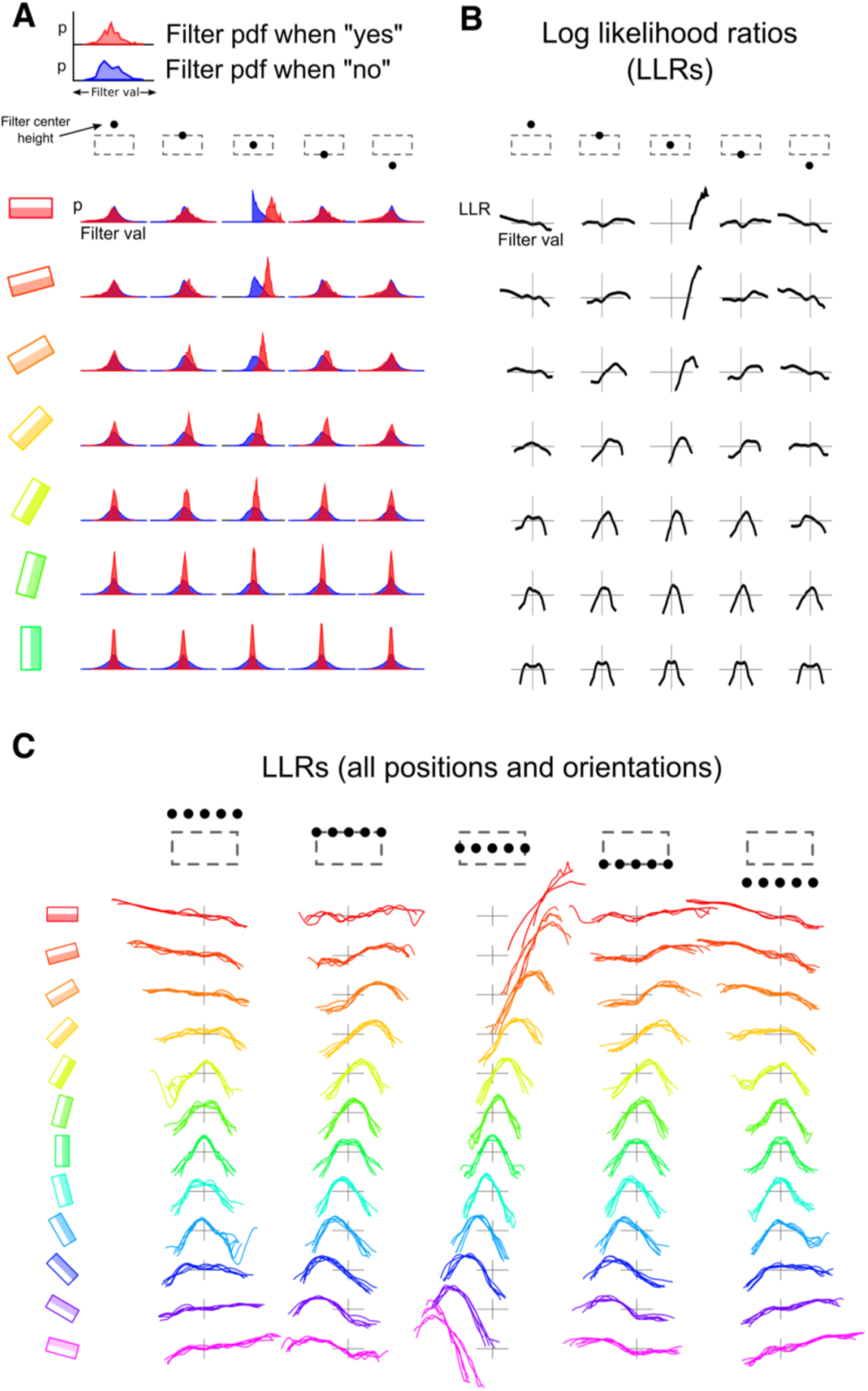
Computing LLR functions from natural images. **A**, Simple cell responses from 30,000 labeled image patches potentially containing boundaries at the reference location (dashed box) were separately histogrammed for “yes” (red) and “no” (blue) cases. Yes (no) cases were those with confidence scores of 4 and 5 (1 and 2). A subset of simple cell response histograms is shown for 7 orientations and 5 vertical positions (centered horizontally). **B,** By dividing the yes and no distributions and taking natural logarithms, one obtains the LLR functions *λ*_*i*_() for each cell which vary as a function of the cell’s response *r*_*i*_. (**C**) The full set of 300 LLR functions reveals a regular pattern over orientation and location. Cases grouped within each subplot are for 5 horizontal shifts (indicated by black dots at top). Many LLR functions are non-monotonic functions of the cell’s response.

**Figure 4.**
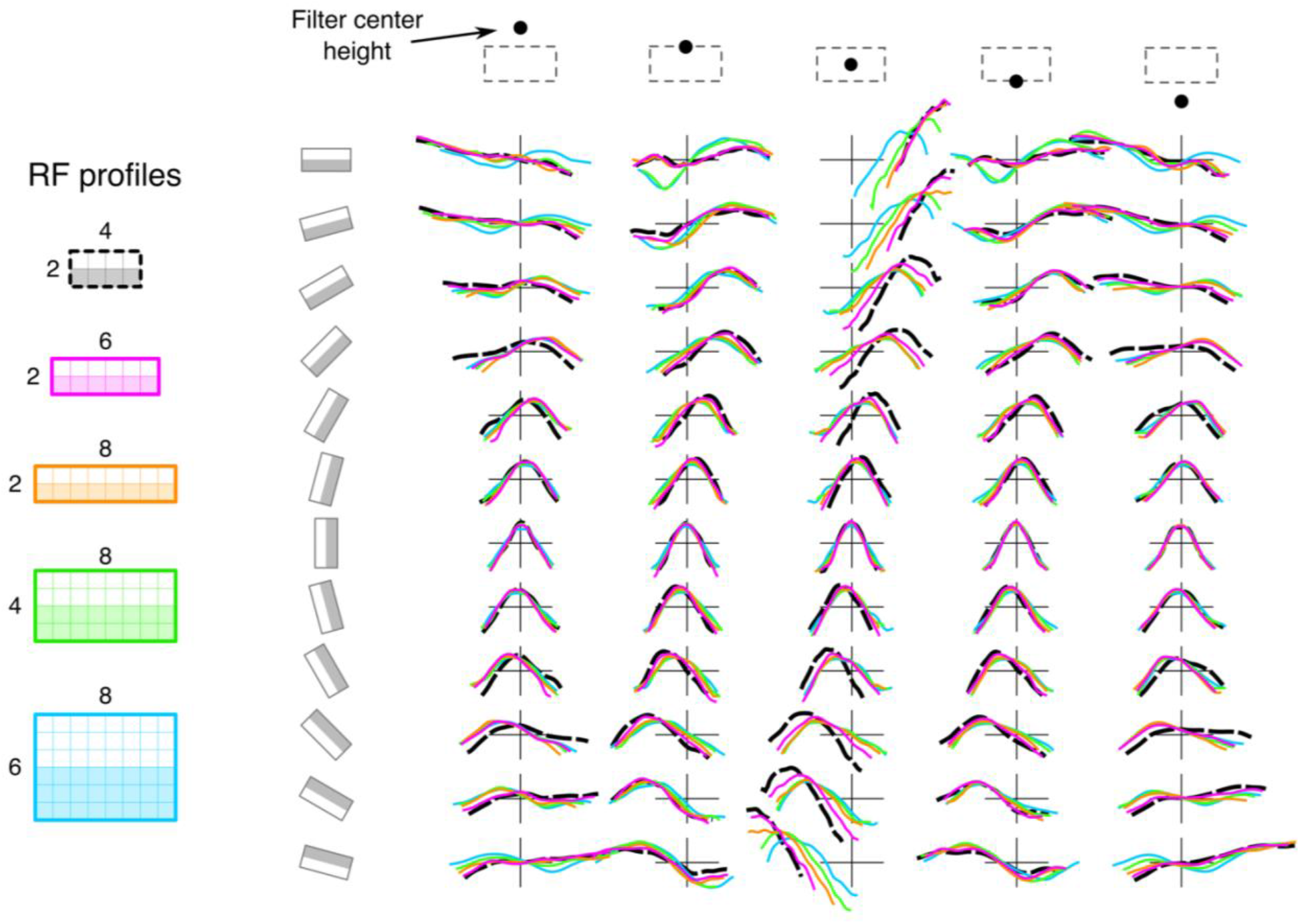
The basic pattern of LLRs forms is conserved across different filter spatial profiles. LLRs were generated for each of the filter profiles shown on the left (2×6, 2×8, 4×8, and 6×8 pixels). The overall spectrum of LLR shapes remains similar for the different cases.

**Figure 5.**
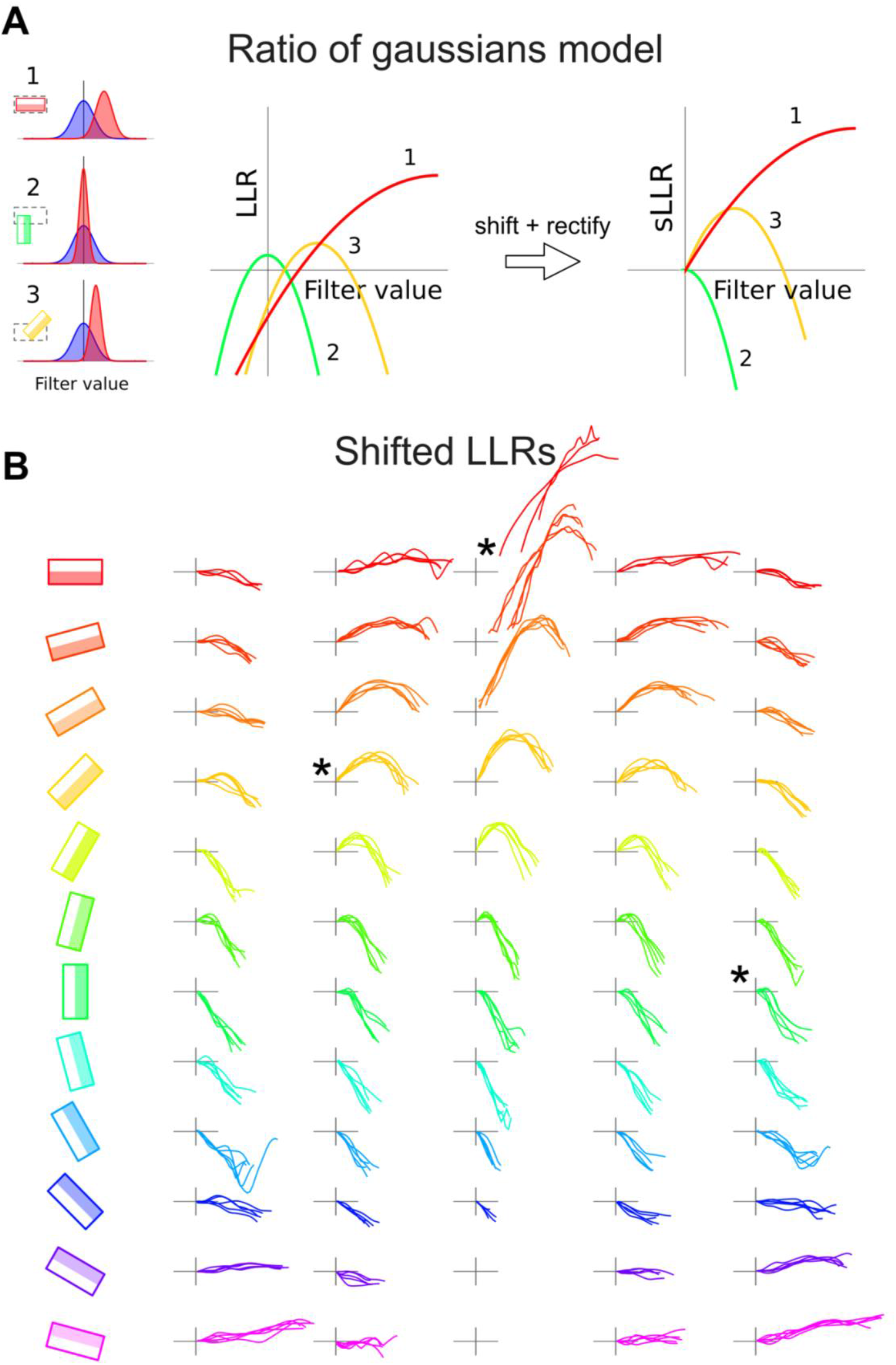
Interpreting the LLRs as cell-cell interaction functions. **A**, Modelling the “yes” and “no” distributions as gaussians (left panel) leads to parabolic LLRs (middle panel). In order to interpret the LLRs as cell-cell interactions functions, we perform two additional processing steps: (1) When a simple cell is inactive, it should not influence the boundary cell; this is accomplished by shifting the LLR to have zero output (y=0) when the input is zero (x=0); sLLR stands for “shifted LLR”. (2) Simple cells cannot have negative firing rates, and so the left halves of the LLRs, corresponding to negative simple cell firing rates, are discarded (these cases are handled by an opponent SC whose RF is identical but with the ON and OFF subfields reversed). This produces the curves in the right panel. **B**. The full set of LLR interactions processed in this way. Many of them are non-monotonic, indicating that that simple cell should have a non-monotonic effect on the boundary cell. The plots corresponding to the three LLRs modelled in panel A are marked with asterisks.

**Figure 6.**
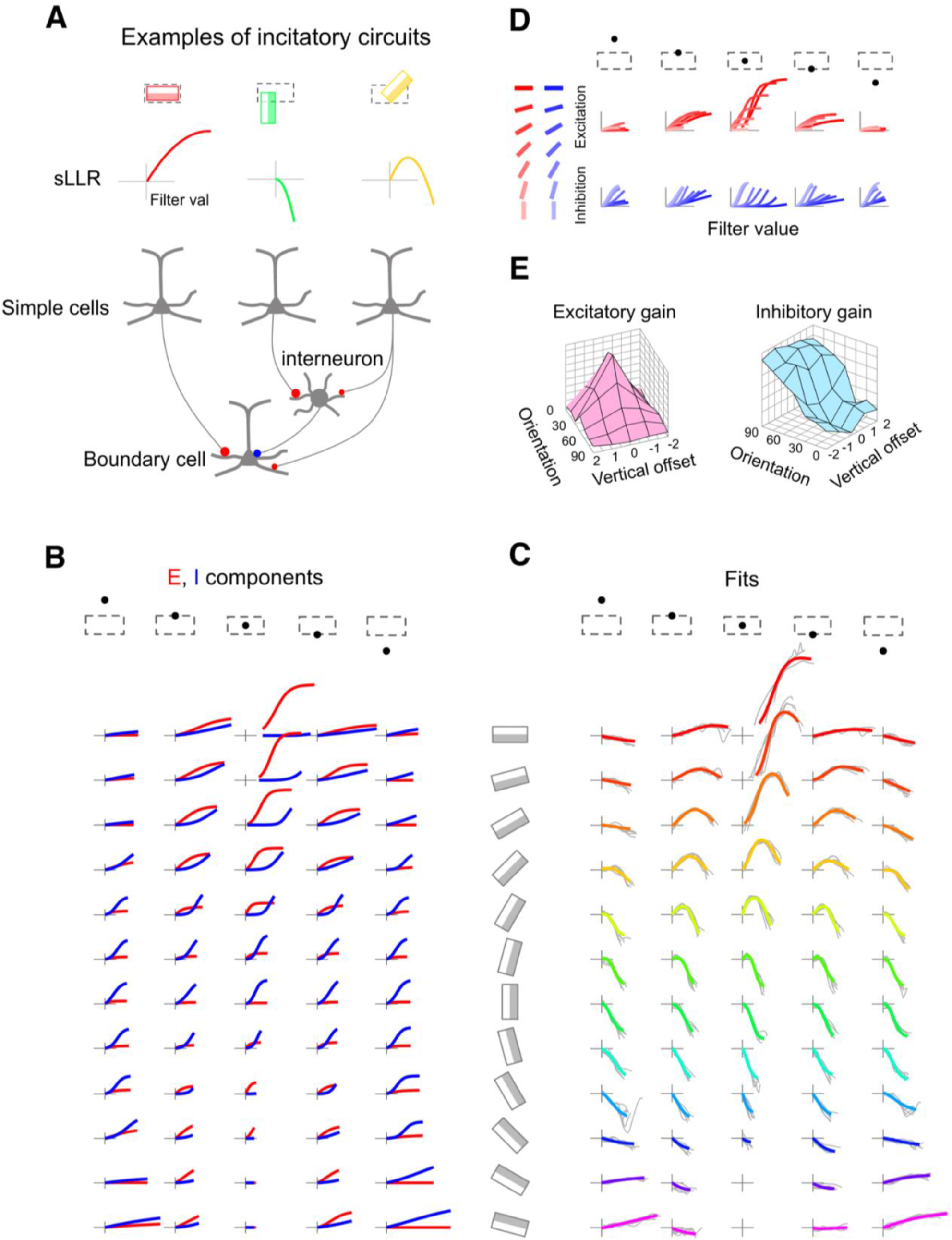
Fitting simple cell–boundary cell interactions (LLRs) with a difference of sigmoids representing separate E and I effects. **A**, Each of the three sLLRs shown can be parametrized by an incitatory circuit. The circuits implementing the red and green sLLRs involve pure excitation and inhibition, special cases of incitation, while that of the orange sLLR involves a nontrivial combination of both excitation and inhibition. **B**, E (red) and I (blue) sigmoidal curves were optimized by manipulating their thresholds, slopes and asymptotes so that their difference fit the corresponding LLR shown in panel **C**. **C**. LLR fits are shown in color, on top of the 5-curve groups from Figure 3C shown in light grey. **D**, E and I sigmoids from b are collected across orientations within each subplot, showing smooth progressions of sigmoid parameters. **E**, Plots show gains for the E and I interaction components. For groups of simple cells horizontally centered at the RL, excitation delivered to the boundary cell becomes weaker and inhibition grows stronger as the neighbor’s orientation deviates from the reference orientation.

**Figure 7.**
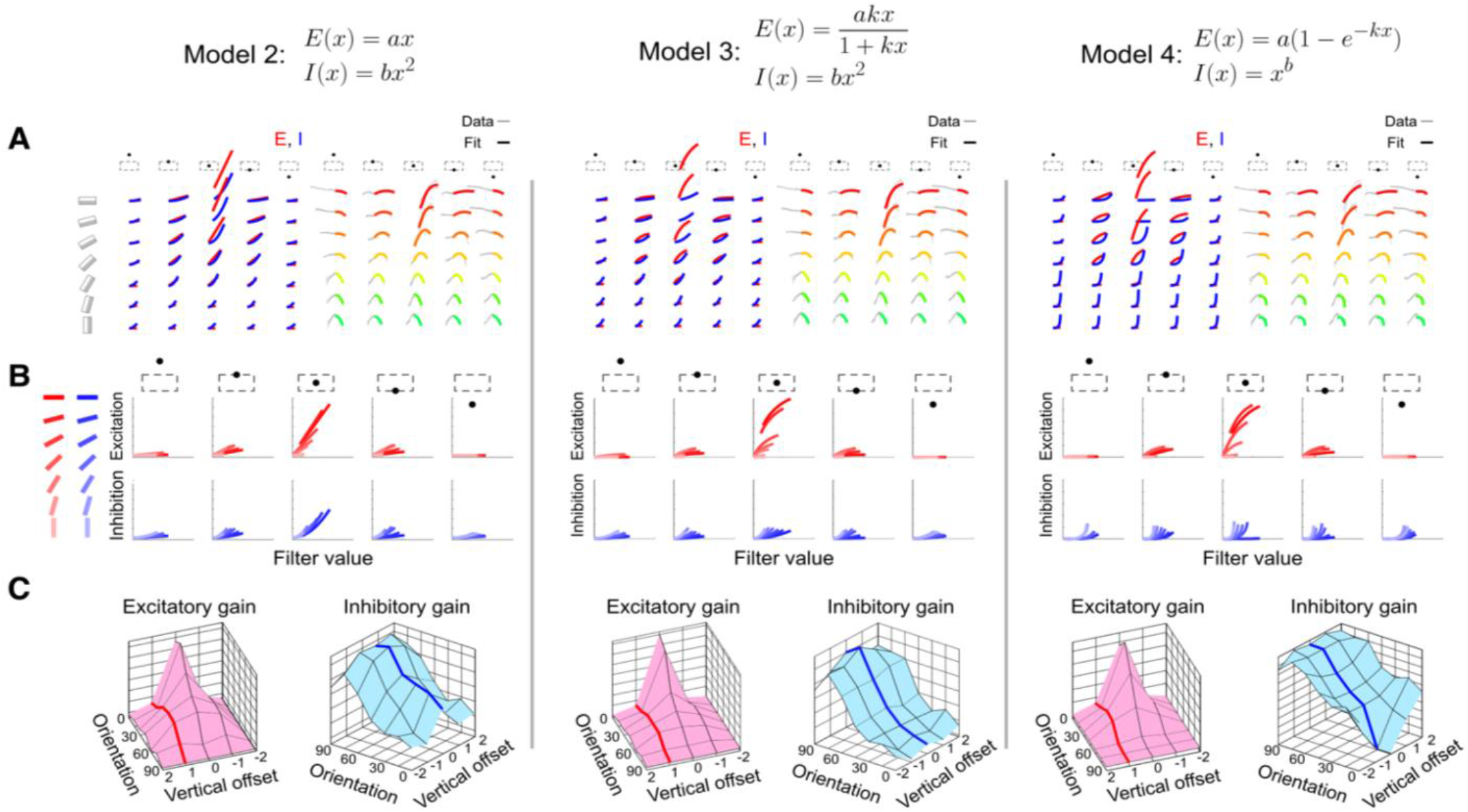
Circuit-level predictions depend only weakly on the choice of parameters representing the excitatory and inhibitory component curves. Three roughly similarly performing models are shown. **A**, Excitatory (red) and inhibitory (blue) curve components (left) and resulting LLR fit (right) are shown for each model. Fit quality is comparable across all three models, and the original model shown in Figure 6C. **B**, Despite having different E-I curve shapes, all three models show the same basic trends in the progression of excitation and inhibition as a function of orientation and vertical offset from the RL. **C**, Summarizing each E and I curve with a single gain parameter shows a similar pattern for the three models.

The same procedure was repeated using different simple cell profiles (2×6, 2×8, 4×8, and 6×8 pixels) to generate the LLR functions shown in Figure 4.

### Modeling log likelihood ratio functions λ(*r*) as differences of sigmoids

As a step in the direction of a circuit level model, we fit the measured LLR functions λ(*r*) for all 300 cells surrounding the reference location with a difference of two sigmoids, 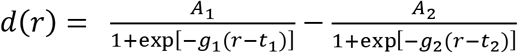. For each cell, approximate amplitudes *A*, gains *g*, and thresholds *t* for the two sigmoids were chosen automatically by minimizing mean-squared error (MSE). The parameters of the sigmoids were then finely adjusted by hand so that the difference of the two sigmoids visually matched the measured λ(*r*) function as closely as possible. We found visually-guided fine optimization better captured the essential shape structure of the LLR function compared to conventional quantitative measures such as MSE. A similar fitting procedure was used for the three models in Figure 7 (model details can be found in the figure and caption). The risk that human visually-guided optimization of curve shape would alter our conclusions was minimal since (1) human visually-guided optimization is based on a much more sophisticated shape-based metric than, say, MSE, and can therefore be reasonably considered as “ground truth”; (2) our conclusions do not depend on quantitative comparisons of fit quality for different models; and (3) the ability to precisely match individual LLR function shapes using a difference of two simple functions is mainly of didactic interest; the more practically significant question is whether a weighted sum of simple excitatory and inhibitory functions (which will in general involve more than two curves) can produce the LLR-like interactions needed to produce improved boundary detection performance (see Figure 8).

**Figure 8.**
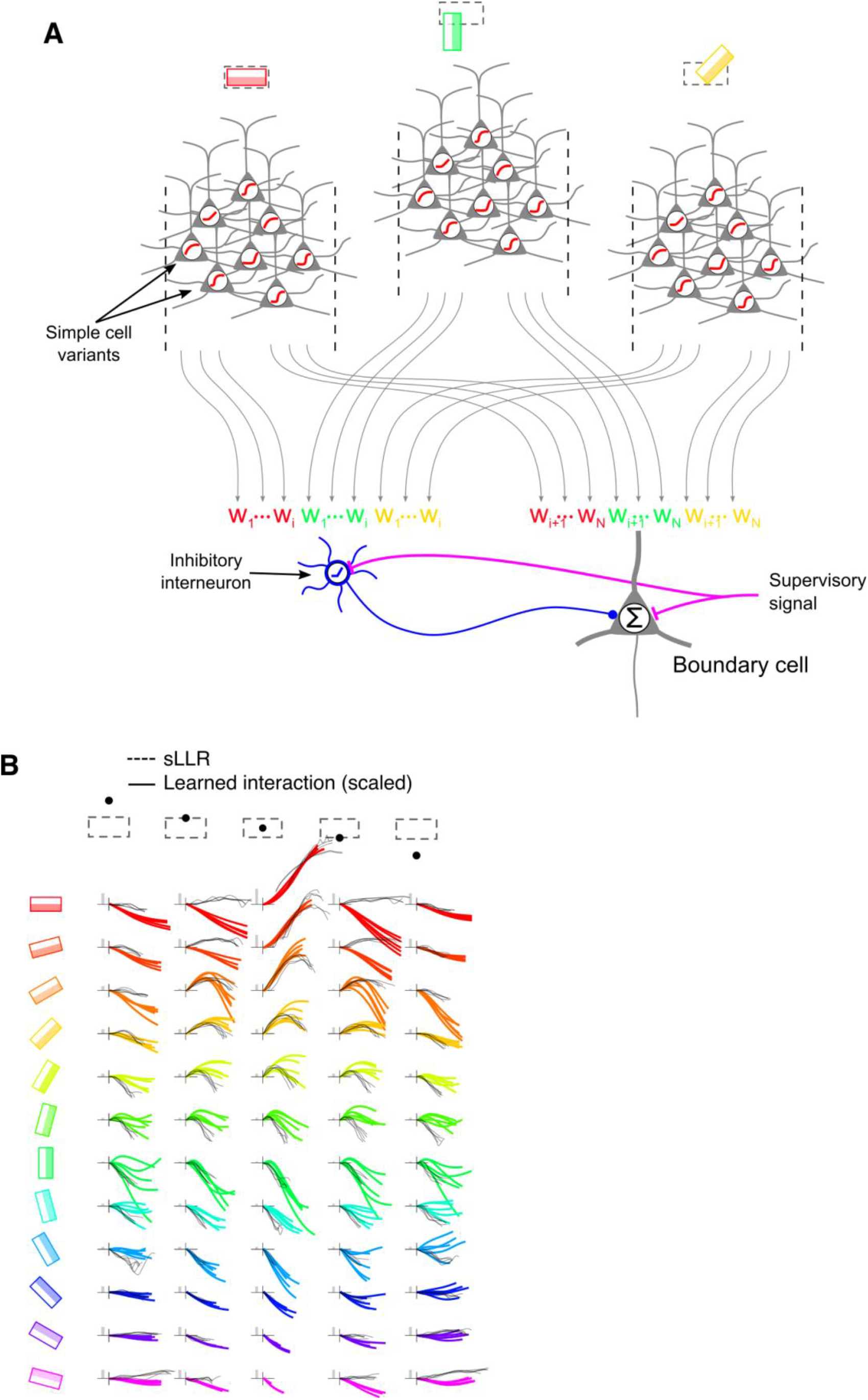
Simple cell-boundary cell interactions can be learned by a biologically plausible synaptic plasticity rule. **A**, Each oriented filter was represented by a population of 8 simple cells, each with a different fixed i/o nonlinearity. Nonlinearities were sigmoids, 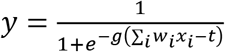, with threshold *t* set at 8 evenly spaced values between −6 and 35. The learning rule used to adjust the weights from each simple cell onto the inhibitory and boundary cell was: *Δw*_*i*_ = ±*η*(*t* − *y*)*x*_*i*_, where *t* is the “training signal” (1 for boundary, 0 for no boundary), *y* is the response of the boundary cell, *x*_*i*_ is the response of the *i*^*th*^ simple cell, η is the learning rate, and the positive (negative) sign was used for the boundary (inhibitory) cell. In the context of our model, this learning rule is mathematically equivalent (up to a transient initial difference in the learning rate parameter η) to a learning rule which constrains all weights to be positive. **B**, To determine the net effect of each filter on the boundary cell (for comparison to the LLRs), the underlying linear filter value was increased from 0 to 1 while holding all other inputs constant, and the weighted sum of the 8 associated simple cells was plotted (colored curves). Black curves are sLLRs from Figure 5B. The gray bar in each plot represents the weight that the BC puts on that group of 5 colored curves.

For the surfaces in Figure 6E, and Figure 7, excitatory gain was computed by measuring the excitatory component’s average slope between r = 0 and r = 10. Inhibitory gain in Figure 5 was computed in the same way. In Figure 7, each of the inhibition families had only a single parameter; this parameter is what is plotted in the inhibitory gain surface plots.

### Optimizing cell-cell interactions in a cortically-inspired network using gradient descent learning

In the data analysis portions of the paper, the term “simple cell responses” referred to the value obtained by computing the dot product between the receptive field kernel and the underlying intensity image. This operation yielded the simple cell’s linear response value *r*_*i*_ for that receptive field location. In a step towards greater biological realism, for the network-level simulation of Figure 8, each of the 300 oriented receptive fields surrounding the reference location was represented a population of 8 model simple cells, all sharing the same receptive field and thus the same linear response value *r*, but having different output nonlinearities (reflecting natural variability within the neuronal population). The differentiation of responses within each 8-cell subpopulation was achieved by modifying the threshold in the sigmoidal output function applied to each cell’s linear response, 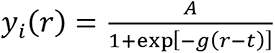. All cells shared the same amplitude and gain parameters (*A* = 1, *g* =0.5), but spanned a range of thresholds from *t* = −6 to 35 in even steps.

The synaptic weights connecting each model simple cell to the boundary cell were obtained as follows. Each image patch created a pattern of activation across the 2,400 model simple cells (300 linear RFs x 8 output threshold variants). We used logistic regression to train a linear classifier to distinguish boundary from non-boundary image patches using the 2,400 model simple cells as inputs. A subset of the data (25,000 of the ∼30,000 labeled patches) was used for training. During training, data was balanced by duplicating boundary-containing patches such that boundary and non-boundary exemplars were equal in number. Training was done using batch gradient descent with a learning rate of η = 0.1, performed for 1000 iterations. The net effect on the boundary cell of the 8 model simple cells sharing the same RF location is visualized in Figure 8B. Each graph shows the weighted sum of the outputs of the 8 simple cell variants as a function of *r*, the cells’ shared underlying linear response value. To facilitate comparison of these graphs with the explicit LLR function *λ*_*i*_(*r*) for that RF location, we scaled the colored interaction functions within each plot. Each plot has one scaling factor that applies to all 5 colored curves in the plot. The inverse of the scaling factor, which can be thought of as the weight that the classifier puts on the curves contained in the subplot, is shown by grey bars.

### Precision-Recall curves measure boundary classification performance

Precision-Recall (PR) curves were generated for the boundary cell network with optimized weights, as well as for the naïve Bayes classifier (based on a literal sum of all cell LLRs; Figure 1BC) and other classifier variants (Figure 8). The term “classifier” refers to any cell or network or mathematical formula that produces a value in response to an image patch, where larger values are meant to signify higher boundary probability. A classifier consisting of the single simple cell at the reference location provided the PR baseline (Figure 8C, blue curve). To generate a PR curve, a classifier was applied to each of the 5,000 labeled but un-trained “test” image patches, and the patches were sorted by the classifier’s output. A threshold was set at the lowest classifier output obtained over the entire test set, and was systematically increased until the highest output in the test set was reached. For every possible threshold, above-threshold image patches were called putative boundaries and below-threshold patches were called putative non-boundaries. “Precision” was calculated by asking what fraction of patches identified as putative boundaries were true boundaries (according to the human ground truth labels), and “Recall” was calculated by asking what fraction of true boundaries were identified as putative boundaries. As the threshold increased, the Precision and Recall values swept out a curve in PR space. Perfect performance would mean 100% Precision and Recall simultaneously, corresponding to the top right corner of the PR graph. Precision-Recall curves are an alternative to ROC curves, and are preferable in domains where the classes are very unbalanced in terms of prior probability. This is the case in our study: boundary images made up only 2.4% of the overall dataset, vs. 97.6% non-boundary cases. See this excellent article for a discussion of this issue (https://machinelearningmastery.com/roc-curves-and-precision-recall-curves-for-classification-in-python/).

### Boundary cell stimulus responses

The idealized boundary image, analogous to a spike-triggered average stimulus, was computed by averaging all natural image patches weighted by their boundary cell response (Figure 11A). For Figure 11B,C, sinusoidal grating stimuli were generated on a 20×20 pixel grid at 24 orientations in 15° steps, and at 60 phase-shifts per cycle. The spatial frequency was chosen to be 0.25 cycles/pixel because it led to relatively artifact-free stimuli at 20×20 pixel resolution, and evoked robust boundary cell responses. For consistency with earlier results, the contrast of each grating image was adjusted to have the same normalizer value (200) as the natural image patches used in the LLR analysis. This was done by generating the grating patch at 100% contrast, computing the normalizer value N on the generated patch using the 100-cell normalization pool (see section above on normalization for details), and scaling down the grating patch by the factor N/200. This procedure reduced the worry that boundary cell responses to the artificially grating stimuli would be distorted by floor or ceiling effects.

**Figure 9.**
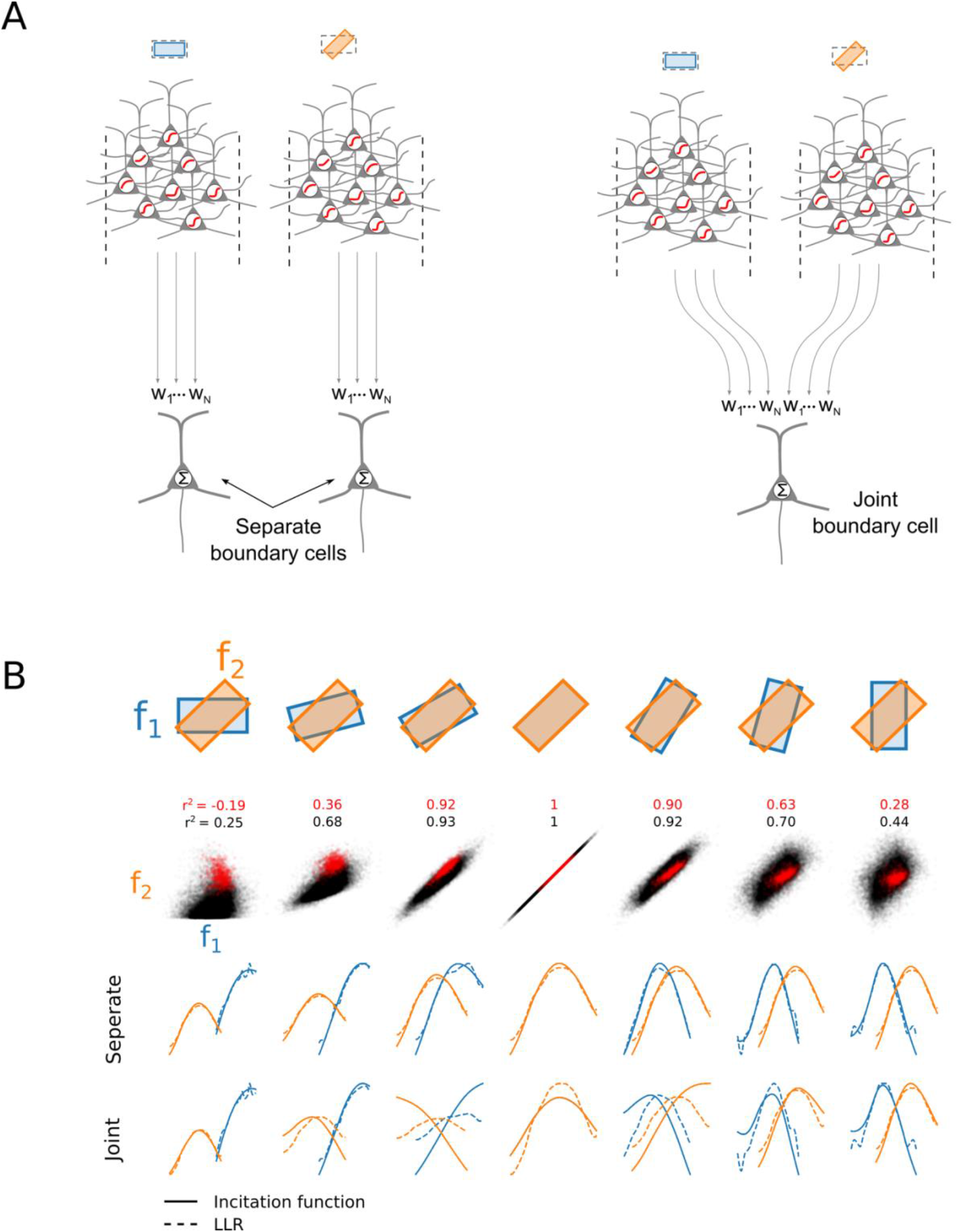
The incitation circuit learns literal LLRs when the filters are class-conditionally independent. **A**, We selected several pairs of filters and fitted either 2 separate incitation circuits, one for each filter (left), or one circuit with both inputs (right). **B**, (Top) Filter pairs ranged from very different (left and right) to very similar or identical (middle) filters. (Middle) Scatter plots of joint filter responses for boundary (red) and non-boundary (black) image patches. (Bottom) When filters were fit separately, the learned incitation functions (solid curves) were nearly identical to the filters’ LLR curves (dashed). When the filters were fit jointly, pairs with very similar filters no longer learned LLR functions due to a breakdown of CCI.

**Figure 10.**
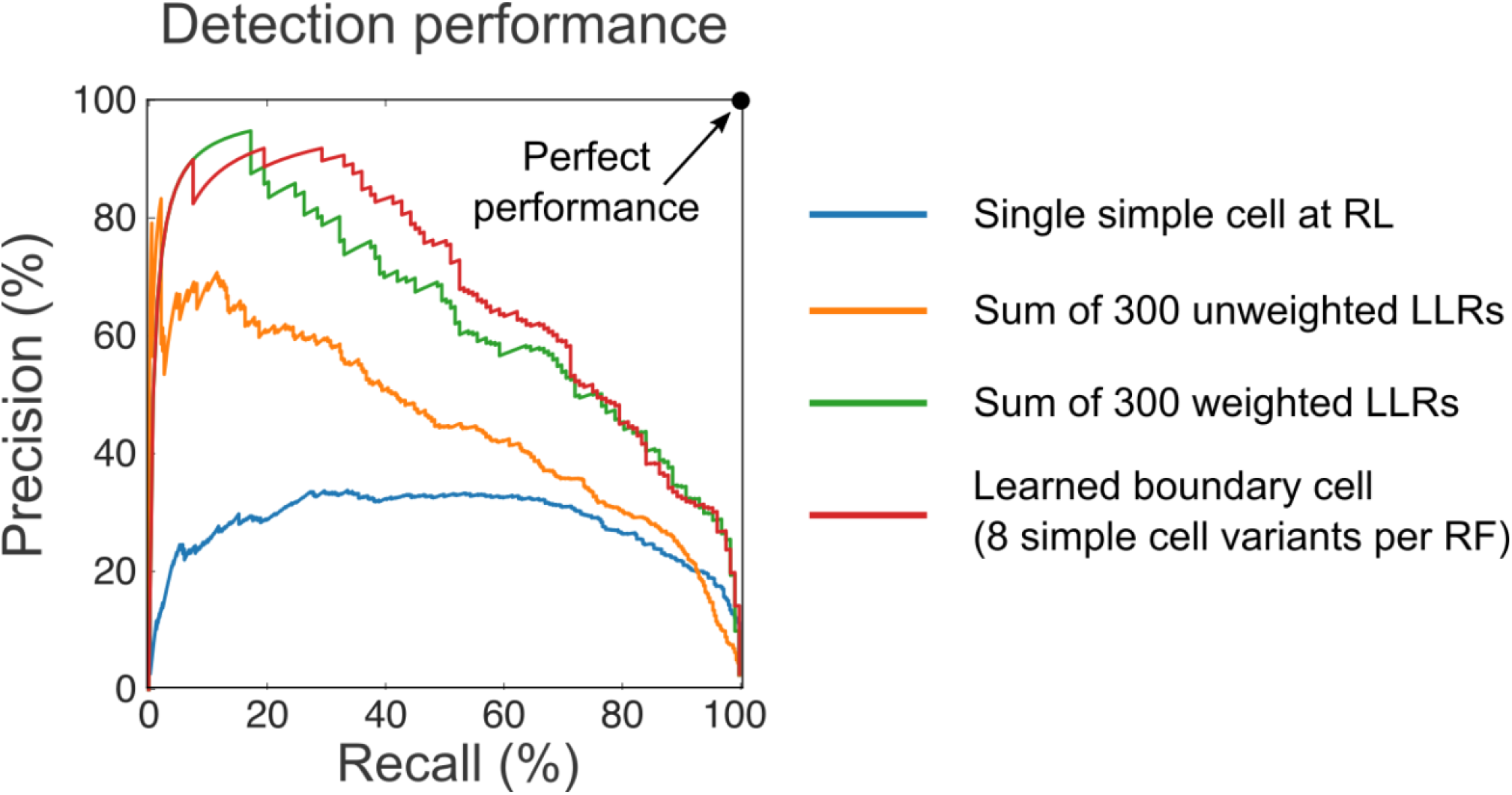
Precision-recall curves for 4 boundary detecting classifiers. See Methods for definitions of precision and recall. Perfect performance is at upper right-hand corner. A single oriented simple cell at the reference location performs poorly (∼20% precision at 90% recall; blue curve). Classifier consisting of a sum of 300 unweighted simple cell LLRs is shown in orange. This case corresponds to the simplified Bayesian classifier depicted in Figure 1C and D. Poor performance was expected given violations of the CCI assumption in the simple cell population. Green curve shows is similar, but with weights on each of the 300 LLRs trained by logistic regression. Performance is dramatically improved, indicating that the learning rule helps to compensate for correlations in the simple cell population. Red curve shows performance of the trained incitation network of Figure 8A. Precision exceeds that of a single simple cell by ∼1.5x at 90% recall, and 2.3x at 50% recall.

**Figure 11.**
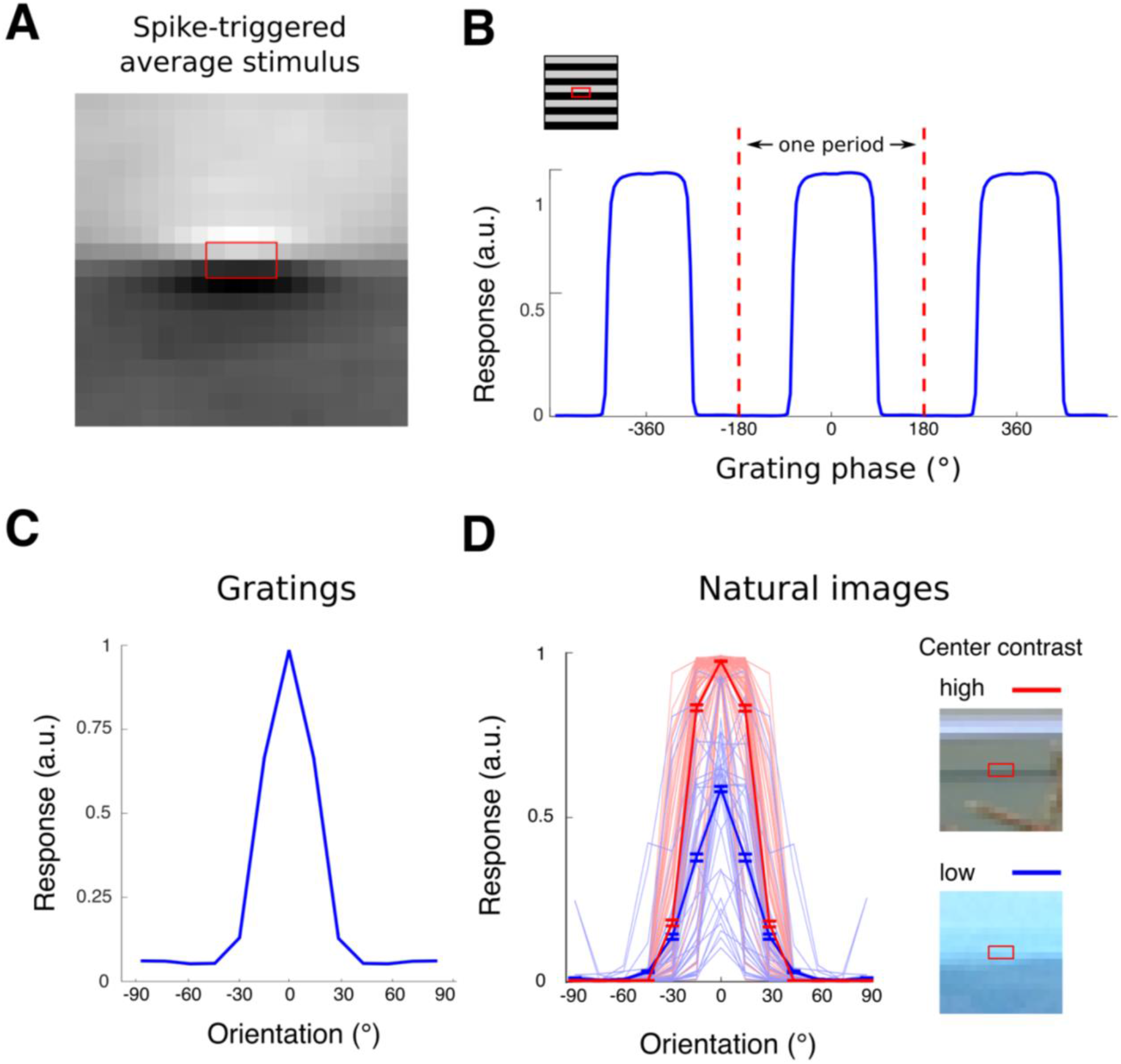
Boundary cell responses to parametric and natural stimuli resemble simple cell responses. To compute BC responses, the weighted sum of LLRs model (orange PR curve in Figure 10) was used. **A**, Spike-triggered average stimulus computed by averaging natural image stimuli weighted by their evoked boundary cell response. **B**, Response of a boundary cell to a grating presented at different phrases. The boundary cell is simple cell-like in that it is sensitive to polarity, responding to only half of all phases. **C**, Orientation tuning curve to the same grating. At each orientation, responses were averaged over all phases of the grating. The resulting tuning curve is similar to those obtained for simple cells in V1. **D**, Patches with fixed surround contrast (normalizer value) and varying center contrast were selected and presented at 15° increments to the boundary cell. For a fixed surround contrast, center contrast increases cell response without increasing tuning width, a hallmark of contrast invariant orientation tuning found in V1 simple cells (full width at half height for high contrast stimuli (red curve) is 43.6°, and for low contrast stimuli, 39.2°).

Simple cell responses to the grating stimuli were presented to the network of Figure 8 just as was done with natural images patches (see above). For Figure 11C, responses were averaged over all phases of the grating at each orientation. Tuning curves in Figure 11D were obtained by presenting natural image stimuli from the N=200 set of normalized image patches. Red and blue curves are for images with 90^th^ and 10^th^ percentile contrast at the reference location, respectively. These percentiles varied in their contrast by roughly a factor of 2. Contrast was defined as the linear filter response at the reference location divided by the average intensity over the reference filter’s 2×4 pixel region of support.

## Results

To gain insight into the cell-cell interactions needed for natural boundary detection, we collected and labeled 30,000 natural image patches, with scores ranging from 5, indicating high confidence that a boundary was present at a “reference location” (RL, indicated by a dashed box in Figure 1A), down to 1, indicating high confidence that a boundary was *not* present at the RL. From these labeled patches, we histogrammed oriented linear filter values (representing simple cell responses) separately for “yes” (scores of 4-5) and “no” (scores of 1-2) cases (red and blue histograms in Figure 3A, respectively). From the responses of 300 neighboring simple cells at 12 orientations on a 5×5 pixel lattice centered on the RL, we computed the likelihoods *p*(*r*_*i*_ | *yes*) and *p*(*r*_*i*_ | *no*), meaning the probability of the *i*^*th*^ simple cell having a particular response *r*_*i*_ when the patch either does (“yes”) or does not (“no”) contain a horizontal boundary. We show in the Methods section that, for a boundary cell to compute the probability of a boundary, and contingent on the assumption that the different filter responses are class conditionally independent, the boundary cell should have as its input the sum of the log likelihood ratio (LLR) functions, 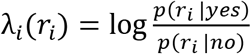 of all contributing simple cells. Each function λ*_i_* (*r_i_*) represents the evidence – positive or negative – that simple cell *i*, when responding at level *r*_*i*_, provides about the existence of a boundary at the reference location. This evidence should be summed by the boundary cell, and then passed through a sigmoidal output function to yield an actual boundary probability (Figure 1C, D).

Accordingly, we computed the LLR functions for all of the 300 simple cells surrounding the reference location. Examples of LLR functions are shown in Figure 3B, and the full set is shown in Figure 3C grouped across 5 horizontal shifts at each orientation and vertical position. The LLR functions varied considerably with position and orientation relative to the reference location, but nonetheless conformed to a small number of qualitative shape prototypes (rising, falling, and bump-shaped). When we generated LLR functions for simple cells receptive fields of different sizes and aspect ratios (2×6, 4×6, 4×8, and 6×8 pixel RF profiles) we found a qualitatively similar pattern of results, indicating that the basic shapes of the LLR functions do not depend sensitively on the RF profiles of the simple cell receptive fields (Figure 4).

The LLR functions λ_*i*_(*r*_*i*_) are important functions to understand since they encode the way a simple cell’s response *r*_*i*_ individually affects boundary probability at a neighboring receptive field location (positively, negatively, or a combination of both). To gain insight into the forms of the λ_*i*_(*r*_*i*_) functions we observed, we developed a simple mathematical model of the process of LLR function formation. If each cell’s yes and no distributions are approximated as gaussian (i.e. unimodal bell-shaped curves) with different means and variances, the resulting LLR functions will be quadratic in form, that is, they will have parabolic shapes. Since the no response distribution is virtually always more variable than the yes distribution, the LLR functions will take the form of downward-pointing parabolas (Figure 5A), qualitatively resembling the LLR functions seen in Figure 3. The particular height and width of each LLR function is determined by the means and variances of the yes and no distributions for that cell (Figure 5A, different colored curves). In addition to qualitatively capturing the range of observed LLR function shapes, this model has a simple interpretation in terms of natural image statistics: in image patches that do not contain boundaries at the reference location (which is to say most image patches), the responses of simple cells in the neighborhood tend to vary widely. On the other hand, image patches that do contain boundaries at the reference location are more constrained, and the responses of nearby simple cells tends to be clustered more tightly around characteristic values, 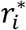. The bump-shaped LLR functions, peaked at 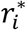, simply reflect the fact that a simple cell provides the strongest positive evidence for a boundary when its response is more typical of “yes” image patches than “no” patches.

To facilitate the interpretation of the λ functions as cell-cell interactions, we slightly reformatted them, in two ways. First, the λ functions were shifted vertically in order that they passed through the origin, reflecting the idea that when a simple cell is not firing (corresponding to *r*_*i*_ = 0 on the graph), its influence on the boundary cell (y-value on the graph) should also be zero. This shift was justified given that the outputs of these functions would later be combined additively (Figure 1C, D), and thus the vertical offsets across the entire population of simple cells could be collapsed into a single net offset at the level of the boundary cell (that would likely be small due to cancellation of positive and negative shifts). Second, simple cell firing rates can only be positive, so the left half of each LLR function, corresponding to a negative simple cell firing rate, was “rectified” (i.e. set to zero). Information was not lost since the same or very similar function would be covered by a different simple cell with the same RF but opposite contrast polarity. The right panel of Figure 5A shows the combined effect of the shift and rectify operations. The full set of shifted LLR functions 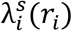 obtained this way is shown in Figure 5B, with the plots corresponding to the conceptual curves in A marked by asterisks.

Returning to the interpretation of shifted LLR functions as simple cell-boundary cell interactions, for some simple cells λ^*s*^ increased monotonically from the origin, meaning that, as the simple cell’s response increased from zero, the evidence it provided to the boundary cell grew steadily more positive. This type of monotonic positive SC-BC interaction was seen for simple cells that were most directly supportive of the hypothesis that a boundary was present at the reference location, such as the simple cell directly overlapping with the RL (middle column in top row of Figure 5B). Referring to the model of Figure 5A, this was a case where the downward-pointing “parabola” peaked far to the right of the origin, so that over the simple cell’s entire observed firing range, its effect on the boundary cell remained on the rising limb of the parabola (case 1 in the right panel of Figure 5A). At even higher firing rates than are plotted in Figure 5B, the λ^*s*^ function would eventually reach its peak and turn back downward, but such high filter values were so rare in yes patches in our natural image data set that the LLR curves could not be reliably estimated beyond the range shown. Two other cases of pure positive λ^*s*^ functions are worth noting: the lower left and right corners of Figure 5B. These cases apply to simple cells whose RFs are nearly “upside down” (i.e. polarity reversed) versions of the reference filter profile, but shifted vertically either 2 pixels above or below the reference location. The fact that these cells are monotonically supportive of the reference hypothesis can be attributed to the existence of many 1-2 pixel wide light and dark horizontal bands in our natural image data set.

For other simple cells, the λ^*s*^ functions *decreased* monotonically from the origin, meaning that, as the simple cell’s firing rate increased from zero, the evidence it provided to the boundary cell grew increasingly more negative. This type of monotonic negative cell-cell interaction was seen for simple cells whose firing supported a hypothesis *incompatible* with the hypothesis that a boundary was present at the RL. The clearest examples of such cells are those with RFs perpendicular to the RL (middle row, green LLR curves. Referring again to the quadratic LLR model of Figure 5A, these monotonically decreasing λ^*s*^ functions arose from cases where the downward-pointing LLR “parabola” was peaked at, or to the left of the origin, so that over the entire response range of the simple cell (to the right of the origin), the λ^*s*^ function fell continuously along its descending limb (as in case 2 in Figure 5A).

For the majority of simple cells, however, the λ^*s*^ functions were bump-shaped, first rising and then falling as the simple cell’s firing rate increased from zero. This type of non-monotonic cell-cell interaction was seen for simple cells whose receptive fields had some degree of overlap in position and orientation with the reference location, so that when a boundary was present at the reference location, their characteristic response levels 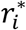 fell in the “middling” range. As a result, these cells provided increasing positive evidence for a boundary at the reference location for response levels up to 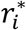, but at even higher response levels, they began to signal stronger evidence for a boundary *at their own RF location and orientation* rather than the reference location. This led to a progressive weakening of the evidence signal sent to the reference boundary cell at response levels above 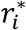.

### A known circuit mechanism can produce the entire observed spectrum of **λ^*s*^** functions

Given that the input to a boundary cell can be approximate as the sum of the λ^*s*^ functions associated with different simple cells (see Figure 1), and that the shifted LLR functions for different simple cells can be either monotonic or nonmonotonic functions of the simple cell’s response, we next asked what kind of neural interconnection circuit is capable of producing the types of monotonic and non-monotonic interaction functions that we observed. One candidate mechanism is the ubiquitous circuit motif in which a cortical cell both directly excites and disynaptically inhibits other cells in its neighborhood (Buzsáki, 1984; George et al., 2011; Isaacson and Scanziani, 2011; Klyachko and Stevens, 2006; McBain and Fisahn, 2001; Pfeffer et al., 2013; Pouille and Scanziani, 2001; Swadlow, 2002; Wehr and Zador, 2003). For monotonic positive or negative SC-BC interactions, the effect could in principle be mediated by pure excitatory or inhibitory connections, respectively (through an interneuron in the case of a pure inhibitory connection) (left and middle cases in Figure 6A). Non-monotonic cell-cell interactions, however, would seem to require a compound excitatory-inhibitory (E-I) interconnection scheme (Figure 6A, rightmost case), wherein the excitatory effect dominates at low firing rates and the inhibitory effect dominates at high firing rates. To determine whether this circuit motif can produce the full range of cell-cell interactions contained in our data set, we assumed that both the direct excitatory and indirect inhibitory pathways exert a sigmoidal effect on the boundary cell, and therefore fit each LLR function with the difference of two sigmoid functions (i.e. assuming the boundary cell sums the excitatory and inhibitory contributions). Each of the sigmoids was allowed to vary in threshold, gain, and amplitude (Figure 6B). The fits are shown in Figure 6C, confirming that the range of cell-cell interactions needed to calculate boundary probability in natural images, including non-monotonic interactions, can be produced by a simple model of a circuit motif known to be present in V1 (see reference list just above). To determine whether the successful fitting of λ^*s*^ functions depended on our particular choice of sigmoidal E and I basis functions, we repeated the fitting procedure using 3 alternative sets of E and I sigmoidal basis functions and obtained similar results (Figure 7). This indicates that the basic types of cell-cell interactions needed to detect object boundaries in natural images can be produced easily by this general type of compound E-I, or “incitatory” circuit.

We next looked for regularities in the progression of excitatory-inhibitory curve pairs used to fit the λ^*s*^ functions depending on a neighbor cell’s offset in position and orientation from the reference location (Figure 6D). We observed the following patterns. First, as the neighbor’s orientation offset from the RL increases and approaches 90 degrees (indicated by lightness changes within each plot), excitation becomes weaker, and inhibition becomes both stronger and lower in threshold, resembling cross-orientation suppression (a staple function of V1 (Bishop et al., 1973; DeAngelis et al., 1992; Geisler and Albrecht, 1992); though see Priebe and Ferster, 2006). Second, we observed a gradual weakening of both excitation and inhibition as a neighbor cell moves further from the RL in the direction perpendicular to the boundary orientation (different plot columns), reflecting the expected decline in informativeness as a neighbor cell moves further from the boundary cell in question. To probe this effect further, we characterized each excitatory and inhibitory curve by its gain parameter and plotted the gains separately as a function of a neighbor’s orientation difference and spatial offset relative to the RL (Figure 6E). These surfaces confirm that, under this simple difference-of-sigmoids model, the strength of the excitation and inhibition imparted to a boundary cell by neighboring simple cells varies systematically with offset in RF position and orientation. The pattern is non-obvious, however, so that if measured experimentally, it could be difficult to interpret without the benefit of a normative framework such as the one we have adopted here.

### Optimizing the parameters for a boundary-detecting incitation network

In pursuit of our goal to understand how cells in V1 detect natural object boundaries, our approach thus far has been to frame boundary detection as a Bayesian classification problem, where the inputs are simple cells (with the simplifying assumption that all simple cells are CCI) (Figure 1); collect ground truth data from human-labeled natural images (Figures 3, 4); and calculate what the simple cell-boundary cell interactions should look like in situations where the CCI assumption holds true. The approach has led to two main findings. First, simple cell-boundary cell interaction functions λ_i_^*s*^ are in general nonlinear and often nonmonotonic (Figure 5), and so cannot be captured by a direct connection from an SC to a BC through a conventional positive or negative weight. On the other hand, we showed that the entire family of rising, falling, and bump-shaped SC-BC interaction functions *can* be captured by a 2-stage “incitation” circuit motif that is known to exist in V1 (Figure 6A, 7).

The precise forms of the SC-BC interaction functions that we might expect to find in V1 remains in question, however: The 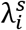 functions shown in Figures 5B and 6C are only the optimal cell-cell interaction functions if all of the simple cells impinging on the boundary cell are CCI. To reiterate, this rather severe condition means that no simple cell provides any information about any other simple cell’s activity level either on or off object boundaries (see Methods and Appendix for details). The condition is, of course, trivially true for a *single* simple cell, and in that case, the simple cell’s functional connection to the boundary cell should be exactly its LLR. The assumption may also hold true for a small population of minimally overlapping simple cells (see Ramachandra and Mel, 2013), but it is definitely not true for 300 simple cells with densely overlapping receptive fields, which is the scenario we consider here. Generally speaking, if the number of simple cells exceeds the number of underlying stimulus dimensions, which is at most 64 in our case (i.e. the number of pixels covered by the 300 simple cells’ RF profiles), then the representation is “overcomplete”, and the simple cells are necessarily correlated. For such a correlated cell population, the likelihood terms in Eq. 2 (pink shading) can no longer be factored, and boundary probability can no longer be expressed as a sum of cell-specific LLR terms λ_*i*_(*r*_*i*_).

This leads to the following interesting question: instead of imposing parameters on the incitation circuit in order to represent cell-specific 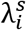 functions as we did in Figures 6 and 7, suppose the synaptic weights of an incitation circuit are trained to maximize boundary detection performance. What would the individual SC-BC interaction functions look like then? Would they be similar to their cell-specific 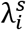 functions? Or would the SC-BC interaction functions be significantly altered, presumably due to correlations within the simple cell population?

To answer this question, we set up a slightly augmented incitation network (Figure 8A) whose modifiable parameters included (i) the excitatory weights connecting each simple cell to the boundary cell, and (ii) the excitatory weights connecting each simple cell to the inhibitory interneuron. In addition, to make it possible for the network to learn nonlinear SC-BC interaction functions by modifying only a single layer of excitatory weights, each simple cell was replicated 8 times to form a small population of closely related cells, all sharing the same oriented receptive field, but each having a different firing threshold. The threshold variability can be seen as arising from natural variation in neuron size, morphology, firing dynamics, etc. The functional purpose of this scheme is that each simple cell’s activation level (before the threshold is applied) is effectively being recoded through a set of fixed nonlinear basis functions, which facilitates learning. The regularly spaced threshold settings used for the groups of 8 cells are given in the Methods section. Each pre-synaptic simple cell acted on the boundary cell through two adjustable weights, one excitatory weight directly onto the boundary cell, and one excitatory weight onto the boundary cell’s inhibitory “partner” cell, which would contribute to disynaptic inhibition of the boundary cell (Figure 8A). The inhibitory neuron was modeled as a linear cell whose firing rate was a weighted sum of its synaptic inputs. Three examples of oriented receptive fields (red, green, and yellow) and their associated simple cell variants are depicted schematically in Figure 8A.

Training occurred as follows. Labeled image patches containing boundaries and non-boundaries (with equalized probability) were presented to the 2,400 (=300×8) simple cells; ground-truth labels from the natural image dataset were presented to the boundary cell (1 for boundary, 0 for no boundary); and the excitatory synapses between the simple cells and the boundary cell and its associated inhibitory neuron were adjusted using a supervised logistic regression learning rule (Murphy, 2012). We then performed virtual neurophysiology to probe the net effect of each oriented receptive field on the boundary cell’s response induced by the 8 simple cell variants (i.e. basis functions) sharing that RF.

The learned interaction functions again included monotonic rising and falling as well as non-monotonic bump-shaped functions (Figure 8B, colored curves). For some cells the learned SB-BC interaction functions corresponded closely to their respective sLLRs (thin grey lines), most notably the cells centered on the RL at all different orientations (middle column of Figure 8B). For this category of cells, the simplified Bayesian formulation seems to explain their role in the boundary detection computation, but no strong inference can be made along these lines given that we could neither predict, nor retroactively account for, which cells fell into this category. In other cases, one or two of the learned SC-BC interaction functions in each group of 5 overlapped heavily with their corresponding sLLR curves, whereas the other curves in the group were driven apart by the learning rule to cover a much wider spread (vertically or horizontally or both) than the original set of sLLRs. In still other cases, the learned interaction functions were nearly “opposite” to their corresponding sLLR functions (e.g. red curves in columns 2 and 4 of the second row in Figure 8B). In these cases, also, we were unable to explain why the cells’ learned interaction functions deviated so extensively from the predictions of the simplified Bayesian model. In the hopes of at least confirming that the deviations were caused by violations in the CCI assumption, we conducted a simple experiment, described next, in which correlations between pairs of simple cell inputs to a boundary cell were systematically manipulated.

### Probing the relationship between the incitation circuit and Bayes rule

To probe the role of correlations between simple cells in shaping SC-BC interaction functions in a boundary-detecting circuit, we ran a simple experiment in which a boundary cell received input from just two simple cells whose RFs overlapped to varying degrees. For each simple cell pair, we fit the parameters of the incitation circuit either separately (Figure 9A, left) or jointly (Figure 9A, right). We tested pairs of filters ranging from very dependent (Figure 9B, middle columns) to nearly independent (Figure 9B; outer columns). Scatter plots of joint filter responses to boundary (red) and non-boundary (black) patches are shown below each pair. When the SC-BC interaction functions were learned separately, they were nearly identical to the literal LLRs (Figure 9B, first row of blue and orange curves; solid curves show learned interactions, dashed curves show LLRs). On the other hand, when the SC-BC interactions were learned jointly, for SC pairs with heavily overlapping RFs, which led to a breakdown of the CCI assumption, the learned interactions differed significantly from the pure LLRs (Figure 9B, middle columns). Consistent with these observations, we show analytically in the Appendix that an incitation circuit like the one shown in Figure 8A will learn LLRs if the input features are CCI. Consequently, any observable differences between the learned incitation functions and the LLRs must be attributable to a breakdown of class-conditional independence.

### Comparing boundary detection performance of four models

The optimization of synaptic weights in the incitation circuit of Figure 8A opens up an additional avenue for validation (or refutation) of our overarching boundary cell hypothesis. Our main premise is that a boundary cell in V1 should be able to significantly improve its boundary detection performance (compared to a single simple cell at the reference location) if it can access, through the local cortical circuit, a large and diverse set of simple cells covering the neighborhood. A major additional claim is that the incitation circuit motif, which is known to exist in V1, is well suited to deliver such a performance improvement. To directly test these claims, we compared the Precision-Recall performance of the trained incitation network (Figure 10, red curve) to 3 other boundary detectors: (1) the “null hypothesis”, consisting of a single conventional simple cell centered at the reference location (blue curve); (2) an unweighted sum of 300 literal LLRs (orange curve); this is essentially a direct implementation of Bayes rule under the CCI assumption (see Figure 1D); and (3) a *weighted* sum of the same 300 literal LLRs (green curve); this hybrid model honors the basic structure of the Bayesian classifier of Figure 1D, but allows weights on each of the LLR inputs to help compensate for CCI violations in the simple cell population.

The results shown in Figure 10 support 3 conclusions: (1) the superior performance of all 3 multi-input classifier variants compared to a single conventional simple cell reinforces the point that individual simple cells are seriously underpowered as natural boundary detectors (see Ramachandra and Mel 2013); (2) the superior performance of the 2 classifier variants with learned synaptic weights (red and green curves) compared to the simplified Bayesian classifier that receives unweighted LLR inputs (orange curve) attests to the importance of a learning rule that is sensitive to natural image statistics and can help compensate for unwanted input correlations; and (3) the very similar performance of the optimized incitation network (red curve) and the weighted sum of LLRs (green curve), especially in the high recall range, is again suggestive of a non-trivial connection between the simplified Bayesian classifier of Figure 1A and the behavior of the learned boundary detecting incitation circuit of Figure 8A.

We conclude by noting that the requirements for developing a cortical circuit that produces significantly improved boundary detection performance compared to a conventional simple cell are relatively modest, including (1) a compound E-I circuit motif that we have dubbed an “incitation” circuit, which is known to exist in V1; (2) variability in firing thresholds across the population of simple cells; and (3) a supervised “delta rule” capable of setting the strengths (and/or dendritic locations) of the excitatory contacts from simple cells onto boundary cells and their associated interneurons. Possible sources of the supervisory signal are taken up in the Discussion.

## DISCUSSION

In the 60 years since Hubel and Wiesel first discovered orientation-tuned simple cells in V1, it has been generally assumed that these cells contribute in some way to the detection of object boundaries (Angelucci et al., 2002; Field et al., 1993; Grosof et al., 1993; Kapadia et al., 1995, 2000; Polat et al., 1998; Sceniak et al., 1999). Consistent with this idea, virtually every modern object recognition system, whether designed by hand or trained from natural image data, includes simple cell-like filtering in its early stages of processing (Fukushima et al., 1983; Krizhevsky et al., 2012; Lades et al., 1993; Lecun et al., 1998; Mel, 1997; Riesenhuber and Poggio, 1999). Surprisingly, however, the quantitative relationship between simple cell responses, typically modeled as divisively normalized linear filters (Carandini and Heeger, 2012), and object boundary probability in natural images, has been little explored (though see Ramachandra and Mel, 2013), making it difficult to know whether or how V1 circuits contribute to this behaviorally relevant natural computation. It is important to emphasize that a simple cell on its own is a poor detector of natural object boundaries within its receptive field (see also Arbelaez et al., 2011): as shown in Figure 10 (blue curve), if we use a simple cell’s response as an indicator of the presence of an object boundary within its RF, even when the threshold for detection is raised to such a high value that half of all true boundaries are rejected (corresponding to a Recall score of 50%), almost two thirds of the “detected” edges at that threshold are false positives (corresponding to a Precision score of ∼35%). The reason a simple cell is such an unreliable edge detector is that true object boundaries are rare (the overwhelming majority of points in Figure 12A are piled in the lower half of the plot), and when they do occur, they are very often of low contrast. Much more common are high contrast non-edge structures (e.g. textures) that contain sufficient oriented energy to strongly drive simple oriented filters.

**Figure 12.**
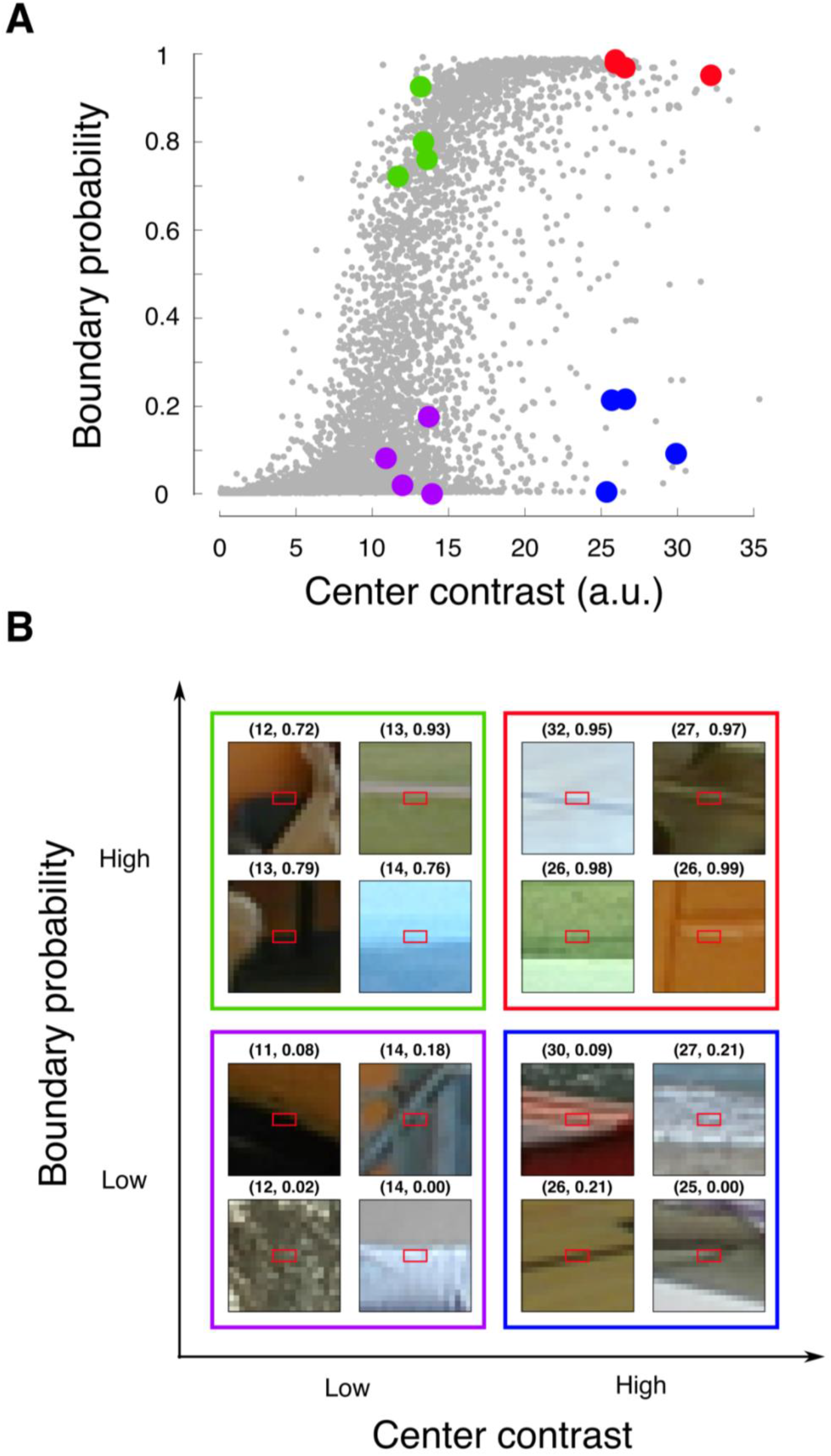
Distinguishing linear filter responses from boundary probability responses. To determine whether a given cell is computing linear contrast or boundary probability, it is necessary to use a stimulus set which dissociates these two measures. Roughly speaking, what is needed are stimuli whose linear filter and boundary probability scores are “well spread” throughout linear filter-boundary probability space. **A**, Plotting the two scores for all labelled patches shows that they are highly correlated, and that randomly selected patches are likely to lie at the lower left and upper right corners of this space – where linear contrast and boundary probability are either both low or high together. Therefore, if only these stimuli were presented to the cell, it would be difficult to know whether high cell responses were being driven by linear contrast or boundary probability. It would be better to present stimuli that are well spread over the space of the two scores (colored dots) so that cell responses to each variable can be assessed separately. **B**, Examples of these stimuli are shown. They include low contrast non-edges (purple cases), high contrast non-edges (blue cases), low contrast edges (green cases), and high contrast edges (red cases).

The poor boundary detection performance of a lone simple cell leads to the conjecture that V1 also contains “smarter” cells that compute boundary probability by combining the responses of multiple simple cells covering a local neighborhood. In a previous study, we suggested that the appropriate strategy for constructing a boundary cell from a local population of simple cells was to (1) select a small set of simple cells (e.g. 6 cells) that were both individually informative and class-conditionally independent (see Methods for discussion of the CCI assumption); (2) evaluate the log-likelihood ratios for each of the participating simple cells, which would tell us the optimal functional interconnections between each simple cell and the boundary cell (according to Bayes rule); and (3) sum the LLRs and apply a fixed sigmoidal nonlinearity to compute boundary probability (Ramachandra and Mel, 2013) (Figure 1C,D). The present study extends that previous work in eight ways: (1) we collected and analyzed individual LLRs for *all* of the simple cells at all orientations covering a 5×5 pixel neighborhood in the vicinity of a boundary cell’s RF (300 cells total); (2) we show that the idealized functional interconnections between simple cells and boundary cells depend systematically on the relative positions and orientations of the simple cell and boundary cell RFs (Figure 3) – but are relatively insensitive to the scale or aspect ratio of the simple cell receptive fields (Figure 4); (3) we developed a simple analytical model (i.e. gaussian likelihoods->quadratic LLRs) that shows how the three seemingly different types of SC-BC interaction functions – rising, falling, and bump-shaped functions – represent different ranges of the same underlying (quadratic) function class (Figure 5); (4) we show that a mixed excitatory-inhibitory, or “incitatory”, circuit motif that is known to exist in V1 is capable of producing the entire spectrum of natural image-derived SC-BC interaction functions (Figures 6,7); (5) we show that the parameters of a boundary-detecting incitation circuit can be learned by adjusting a single layer of excitatory weights (Figure 8A); (7) we show that a learned incitation circuit can improve the precision of boundary detection in the high-recall range by 43% to 121% compared to a conventional simple cell model (Figure 10); and (8) by “reading out” the weights of the learned incitation circuit, we show that the simple cell-boundary cell interaction functions that we would expect to find in the visual cortex are not likely to be verbatim LLRs, but rather, perturbed versions due to class-conditional dependencies among simple cells whose receptive fields overlap heavily with each other (Figures 8B, 9). This could be helpful in interpreting the results of future neurophysiological experiments in V1.

### Experimental predictions

#### Distinguishing boundary cells from conventional simple cells

Having shown that cortical circuitry is capable in principle of producing boundary cells from simple cells using only a single layer of modifiable excitatory weights, it is important to ask how BCs could be detected experimentally, and distinguished from conventional simple cells (or the simple cell-like subunits of complex cells – Hubel and Wiesel, 1962; Movshon et al., 1978; Ohzawa et al., 1997).

To determine how BCs would respond to various stimuli, stimulus patches were scaled to have the same fixed value of the normalizer used in earlier figures, once again reflecting a simple form of divisive normalization (see Methods section *Boundary cell stimulus responses*). We first constructed a canonical stimulus for a boundary cell akin to a spike triggered average by averaging all image patches weighted by their evoked boundary cell response. As expected, the STA stimulus appears as a localized, polarized, oriented boundary segment reminiscent of a simple cell’s receptive field (Figure 11A). We then presented drifting sine wave gratings covering a boundary cell’s “classical receptive field”, leading to the unremarkable phase response and orientation tuning curves shown in Figure 11B, C. Next, we used labeled natural edges with the same normalizer value to explore the effect of increasing center contrast on orientation tuning curve width. (This was not a perfectly controlled experiment because variations in center contrast at a fixed normalizer value would have led to *anti*-variations in surround contrast, but given the filter value at the RF center was only one of 100 filters of many orientations used to compute the normalizer value, this effect was likely small). Subject to this limitation, as shown in Figure 11B, the boundary cell’s tuning width is essentially constant across a roughly 2-fold change in center contrast – the limit of analysis allowed by our labeled database (average tuning curve has full width at half height for high contrast stimuli 43.6°; for low contrast stimuli, 39.2°).

Thus, for oriented edges and gratings presented within the CRF, boundary cells behave similarly to conventional simple cells in that they (1) have phase-dependent responses; (2) are orientation tuned; and (3) have tuning curves whose widths are roughly contrast invariant (Alitto and Usrey, 2004). It is therefore possible that boundary cells exist and have been classified as conventional simple cells in previous experiments using simplified stimuli. Among the multiple types of V1 cells that have been previously described, boundary cells share most in common with double opponent cells, which are orientation tuned, have mostly odd-symmetric receptive field profiles as would be expected for boundary detecting cells (Ringach, 2002), and respond to boundaries whether defined by luminance or color (Johnson et al., 2008).

In future neurophysiological studies, an efficient means of dissociating conventional simple cells, which respond to oriented contrast independent of boundary probability, from putative boundary cells, which respond to boundary probability independent of oriented contrast, would be to use natural image stimuli drawn from the four corners of the oriented contrast – boundary probability space (Figure 12A). Image patches with low oriented contrast and low boundary probability scores (purple dots) tend to contain flat, unstructured image regions; patches with low contrast and high probability (green dots) tend to contain well-structured, faint edges; patches with high contrast but low probability (blue dots) tend to contain contrasty noise or misaligned edges; and regions with high contrast *and* high probability (red dots) are typically well-structured, strong edges (Figure 12B). This factorial stimulus set would make it possible to identify pure simple cells, pure boundary cells, as well as cells of intermediate type.

#### Diverse inputs to the dendrites of boundary cells?

One of our main findings is that a cell in visual cortex whose job is to detect object boundaries can improve its detection performance if it collects input from many simple cells in its vicinity with a diversity of receptive field positions and orientations. Consistent with this, several recent 2-photon calcium imaging studies have mapped the receptive field properties of individual dendritic spines on V1 neurons in mice, ferrets, and monkeys, and have shown that the inputs to a single V1 cell (and often a single dendrite) are quite variable in terms of their receptive field properties, covering a much wider range of preferred orientations and RF positions than might be expected given the target cell’s more sharply tuned response preferences (Jia et al, 2010; Wilson et al. 2016; Iacaruso et al. 2016; Scholl et al. 2017; Ju et al. 2020). These findings do not prove that many or most cells in V1 are boundary cells, only that most V1 cells appear to receive the requisite diversity of inputs from neighboring cells. What remains to be shown is that the excitatory and inhibitory inputs from surrounding cells are properly weighted and balanced by the local incitation circuit, so as to maximize boundary detection performance. The conceptual experiment described next could help to establish whether the local cortical circuit actually functions in this way.

#### A predictable spectrum of SC-BC interactions?

A key feature of the boundary cell hypothesis is that SC-BC interaction functions take on predictable forms, depending primarily on the offsets in position and orientation between the SC and BC receptive fields (Figure 8b). It may be possible to empirically measure those interaction functions by applying stimuli that distinguish simple cells from boundary cells (Figure 12) while imaging V1 neurons in awake animals (e.g. Tang et al. 2018; Ju et al. 2020). The approach would require the ability (1) to identify pairs of nearby simple and boundary cells and to characterize their RFs, and (2) to transiently activate or inactivate identified cells optogenetically. Then, while presenting carefully curated natural image patches to the boundary cell – that also overlap with the simple cell’s RF, the simple cell’s activity level could be perturbed, and the response changes in the boundary cell measured. The direction and magnitude of the change in the boundary cell’s activity could be compared to the prediction of a trained incitation network (Figure 8B). For a particular simple cell, certain image patches will drive the cell to a level on the rising slope of its SC-BC interaction function, so that a boost in the cell’s activity (through optogonetic stimulation) should in turn boost the boundary cell’s activity – and conversely for suppression of the simple cell’s activity. For other image patches, the simple cell will be firing at or beyond the mode of its SC-BC interaction function with respect to a particular boundary cell, so that a boost in that simple cell’s activity will lead to a suppression of the boundary cell’s activity (and conversely if the simple cell’s activity is optogenetically suppressed).

It is worth noting that we cannot assume that the SC-BC interaction functions measured in this way will look exactly like those produced by a particular trained incitation network, since even if the overall idea holds true, the interaction functions depend on the complete set of simple cells providing input to a particular boundary cell, which cannot be known. Nonetheless, by repeating these types of response manipulations for a large number of SC-BC pairs, we can hope to find a basic correspondence between predicted and measured SC-BC interactions, in the sense that the measured interactions should include pure increasing cases, pure decreasing cases and non-monotonic cases, with a systematic dependence on the spatial and orientation offsets of the simple and boundary cells’ RFs (Figures 6-8).

It is also worth noting that boundary cells need not reside in, or only in, V1. Nothing precludes that cells that signal boundary probability, rather than boundary contrast, may be found in higher visual areas.

### Relationship to previous work on natural image statistics

A number of previous studies have attempted to explain receptive field properties of cells in the retina, LGN and primary visual cortex in terms of natural image statistics and principles such as efficient coding, sparse coding, and independent components analysis (Barlow, 1981; Bell and Sejnowski, 1995; Laughlin, 1989; Olshausen and Field, 1996; Schwartz and Simoncelli, 2001; Zhu and Rozell, 2013). These studies have been mainly concerned with neural *representation*, where the goal is fast/accurate information transmission through a noisy channel, and eventually faithful image reconstruction. In contrast, our work is primarily concerned with neural *computation*, where the goal is to transform the image into a more abstract shape representation that is more directly useful for guiding behavior.

From a different perspective and with a different goal, Geisler et al. (2001) collected co-occurrence statistics of pre-detected local boundary elements in natural scenes, with the aim to predict human contour grouping performance. Their measurements on natural images included the probability of finding a second boundary element in the vicinity of a first boundary element, depending on the relative offsets in position and orientation of the two elements, or whether two spatially offset boundary elements were more likely to belong to the same or different object. Sigman et al. (2001) also studied co-occurrence statistics of pre-detected boundary elements, coming to the conclusion that boundary elements in natural scenes tend to lie on common circles. The goal to characterize the spatial distribution of pre-detected boundary elements in natural scenes in both of these studies contrasts with our focus here on the detection problem, that is, the problem of discriminating object boundaries from non-boundaries based on populations of simple cell responses collected from a local neighborhood in an image. Furthermore, all of the grouping statistics collected by Geisler et al. and Sigman et al. were represented as scalar values linking pairs of locations/orientations. In contrast, our natural image analysis produces *functions* linking pairs of locations/orientations, which capture how a given simple cell should influence a nearby boundary cell as a part of a boundary detection computation. Also unlike these previous studies, we use our data to constrain and to benchmark cortical circuit models.

### Non-monotonic cell-cell interactions have been previously reported

One of our findings is that among the different types of local cell-cell interactions needed for object boundary detection in natural images, many cannot be described as “excitatory” or “inhibitory”, nor can they be represented by positive or negative synaptic weights, but are instead U-shaped functions wherein cell 1 might excite cell 2 at low firing rates, reach its peak excitatory effect at intermediate firing rates, and inhibit cell 2 at high firing rates. U-shaped functions of the opposite polarity can also occur (Figure 8B). Should we find the idea surprising that nearby cells in the cortex act on each other non-monotonically?

From one perspective, one might argue that whenever there are excitatory and inhibitory cells wired together in a circuit motif, perhaps we should be surprised if we did *not* find non-monotonic interactions between cells. For example, in the “inhibition-stabilized network” model (Jadi and Sejnowski, 2014; Ozeki et al., 2009), which accounts for a number of V1 cell response properties, “non-binary” interactions between cells would almost certainly be expected to occur. Nevertheless, there has been a historical tendency to think about cell-cell interactions in the cortex as being of a defined polarity, represented by a positive or negative scalar value, and often subject to simple geometric rules. The notion of “surround suppression”, for example, reflects both of these tendencies (Adesnik et al., 2012; Cavanaugh et al., 2002; Schwabe et al., 2010). Even as the geometric constraints governing cell-cell interactions become more intricate, such as where interconnection strength and polarity depend on distance or relative orientation, the simplification that cell-cell interactions have a defined polarity is often still relied upon. For example, K.D. Miller’s models of map development include short range excitation and medium-range inhibition (Miller, 1994); Angelucci and Bressler’s models include near and far suppressive surrounds (Angelucci and Bressloff, 2006); and several studies support the idea that cortical cells affect each other laterally through bowtie-shaped “extension fields” consisting of patterned arrays of positive and negative coefficients (e.g. (Bosking et al., 1997; Field et al., 1993; Geisler et al., 2001; Kapadia et al., 2000; Li, 1999; Sigman et al., 2001). In all of these cases, one neuron’s effect on another neuron is described in terms of its scalar connection “strength”.

Not all functional interconnections that have been described in the cortex fit such descriptions, however. Examples of activity-level-dependent interactions have been reported, where the strength and even polarity of the connection between cells depends on the activity levels of the sending and/or receiving cells. For example, the responses of amplitude-tuned neurons in the auditory cortex grow stronger as the sound pressure level increases up to an optimal intensity level, and then are progressively inhibited as the sound grows louder (Suga and Manabe, 1982); in V1, surround modulation can switch from facilitating to suppressive with increasing center contrast (Ichida et al., 2007; Nauhaus et al., 2009; Polat et al., 1998; Schwabe et al., 2006; Somers et al., 1998); length-tuned neurons respond best to an oriented stimulus up to a certain length, but are then progressively inhibited as the stimulus grows longer (Anderson et al., 2001); and non-monotonic modulatory interactions between a neuron’s classical and extra-classical receptive fields have been reported (Polat et al., 1998). These data, though unaccompanied by normative explanations, do support the idea that the sign and magnitude one neuron’s effect on another can depend not only on the relative position and orientation of their receptive fields (in the case of vision), but also on their relative activity levels.

Our paper represents a fleshing out of this type of effect, and is to our knowledge the first normative theory, parameterized by natural images, that specifies how intracolumnar cell-cell interactions may help solve a specific, biologically relevant *classification* problem. By analyzing natural image data on and off object boundaries, we showed that the local cell-cell interactions needed to solve this classification problem are not capturable by scalar weights, but are in general nonlinear functions that depend on “all of the above” – relative location, relative orientation, and relative activity levels of the sending and receiving cells. We further showed that the SC-BC functional connections needed for boundary detection are easily produced by a compound E-I circuit motif (see Figure 6) that is known to exist in the cortex (Buzsáki, 1984; Isaacson and Scanziani, 2011; Klyachko and Stevens, 2006; McBain and Fisahn, 2001; Pfeffer et al., 2013; Pouille and Scanziani, 2001; Swadlow, 2002; Wehr and Zador, 2003). Finally, we showed that the synaptic weights that control the net effect of an “incitation” motif are easily learned. Future experiments will be needed to establish whether trainable incitation circuits are actually used to help solve the difficult natural classification problems faced by neurons in V1 and other areas of the cortex.

### How could a properly parameterized incitation circuit develop?

A possible extension of this work would be to address the limitation that the incitation circuit we show in Figure 8A was trained by a supervised learning rule (logistic regression), but without our providing a biologically based account for the source of the supervision. The original purpose of the exercise was to test whether an incitation circuit with a single layer of modifiable excitatory weights is *capable* of performing object boundary detection at a level comparable to an explicit Bayesian classifier. We found this to be true (Figure 10), suggesting that this particular Bayesian-inspired algorithm lies within the computational scope of cortical tissue. The demonstration leaves open the question, however, as to where a supervisory signal might come from during visual development that alerts a boundary cell and its inhibitory partner when an object boundary is crossing through its receptive field. One possible source of supervision would be a population of neurons located within the same or different area that have access to different visual cues, such as cells sensitive to motion-defined boundaries. Such cells are found at many levels of the visual system, including the retinas of salamanders (Olveczky et al., 2003); V1, V2, V3 MT and IT in the monkey (Marcar et al., 1995, 2000; Sary et al., 1995; Zeki et al., 2003); and in multiple areas of the human visual cortex (Dupont, 1997; Larsson et al., 2010; Mysore et al., 2006; Zeki et al., 2003). Topographic feedback projections from motion boundary-sensitive cells in these areas to V1 (or locally within V1) could help to instruct boundary cells in V1 so that they may perform based purely on pictorial cues (i.e. when motion signals are unavailable).

### Limitations of the model

The boundary detection computation that we have studied was inspired by Bayes rule, and is underlyingly a feedforward computation whose core operation is a sum of LLR terms (Figure 1C). Our attempt to map this computation onto a simple, cortically plausible circuit is shown in Figure 8A, in which a layer of simple cells with varying output nonlinearities activates both (1) a “layer” of boundary cells (though only one BC is shown); and (2) a layer of inhibitory cells, one per BC (though only one inhibitory cell is shown – the one assigned to the one shown boundary cell). Each inhibitory cell, in turn, acts on its associated boundary cells through a fixed connection. Given that the circuit of Figure 8A is purely feedforward, ignoring (1) local or long-range feedback connections that are known to exist in the neocortex (Angelucci et al., 2017); (2) nonlinear dendritic integration effects that could also contribute to boundary detection (Jin et al., 2022); and (3) all dynamics at the synapse, cell, and circuit levels, it falls far short of a fully elaborated cortical circuit model. Rather, the circuit model of Figure 8A should be viewed as a demonstration that a known cortical circuit motif – the incitation motif – is capable of producing cells that superficially resemble simple cells, but are much better at detecting object boundaries in natural scenes than the standard simple cell model (Heeger 1992). A worthy long-range goal would be to fold the boundary-detection capability of a properly parameterized incitation circuit into a more comprehensive cortical circuit model that addresses a wider range of physiological phenomena (Ozeki et al. 2009; Zhu and Rozell 2013).

## Abbreviations

SC: simple cell
BC: boundary cell
RF: receptive field
RL: reference location
LLR: log-likelihood ratio
sLLR: shifted log-likelihood ratio
CCI: class conditional independence
PR: precision-recall

## Acknowledgments

Funding for this work was provided by NIH/NEI (EY016093).

## Author contributions

BM and CR conceived of the original project. GM and BM collected the natural image data, performed the analysis, developed the models, and wrote the paper.

## Appendix 1: Logistic regression learns LLRs assuming CCI

We are interested in estimating the probability of some event *y*, in this case whether a patch contains a boundary, from input features 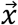, in this case the responses of several simple cells. Logistic regression builds a model of 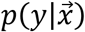 by assuming that the output probability is a sigmoid function of a linear combination of the features:

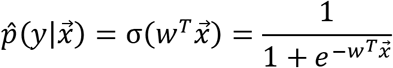

The goal of learning is to pick weights *w* that minimize the expected cross entropy between the true and model probabilities:
1

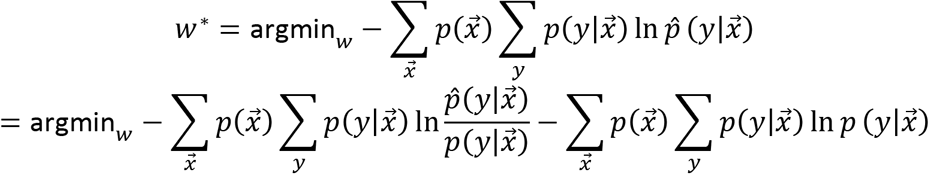

The left term is the KL divergence between the true and model distributions, and the second term is constant with respect to the weights, and can be ignored. The optimization is then

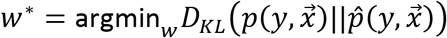

This is minimized when the model distribution *p̂* matches the true distribution *p*. To see that under the assumption of class conditional independence, learning the LLR functions achieves this minimum, observe

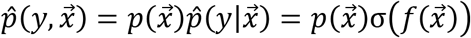

Where 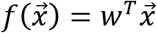. Further, class conditional independence implies

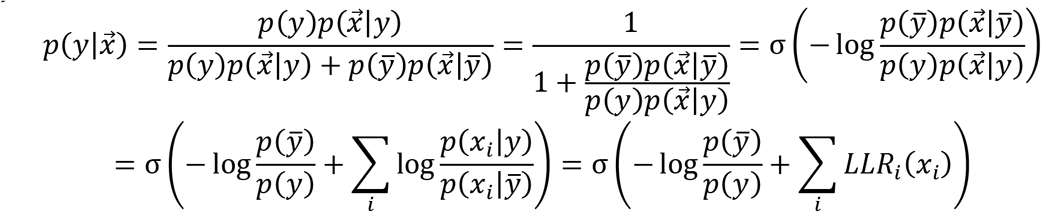

so that the objective can be written

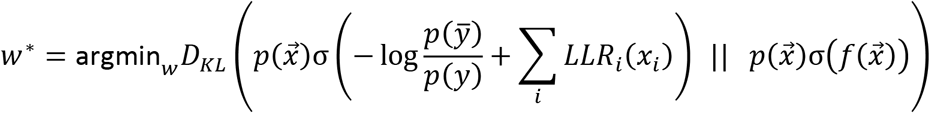

One can see by inspection that the two distributions will be equal and the objective will be minimized if and only if

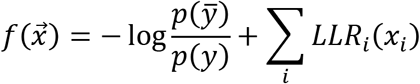

that is, when the classifier simply combines the filter values by passing them through their LLR functions and summing the result.

Now consider the problem of learning the optimal weights through gradient descent using a dataset of N input-output pairs. Call the *i*^*th*^ input patch *x*^(*i*)^ and the *i*^*th*^ output label *y*^(*i*)^. Label non-boundaries *y*_*i*_ = −1 and boundaries *y*_*i*_ = 1. The cost function, gradient, and Hessian can be written

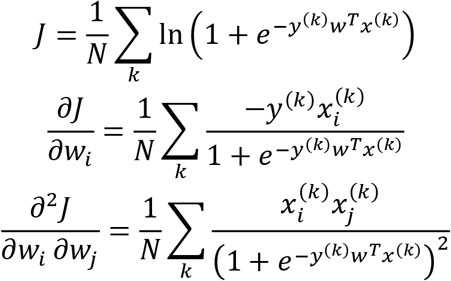

If we collect all of the input patches *x*^(*k*)^ into the rows of the matrix *X*, the Hessian can be written simply as

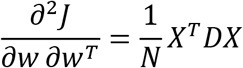

Where *D* is a positive diagonal matrix with entries 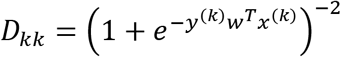. First, note that the Hessian is positive semidefinite since 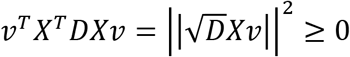 for all *v*. In the case that the *K* features making up the columns of the training set *X* are linearly independent (generically true if the input cells have distinct filter kernels), the Hessian is strictly positive definite since each of the following matrices having rank ≥ *K* implies the same of the next: *X*^*T*^*X*, *XX*^*T*^, *XX*^*T*^*D*, *X*^*T*^*DX*, the last of which is the Hessian. Regardless, by positive semi-definiteness, any local minimum of the objective function *J* is a global minimum, so gradient descent terminates at the global minimum, which by the argument above must be the LLR solution.

